# Haplotype-resolved genome assembly advances genetic understanding of trait diversity in *Phalaenopsis* orchids

**DOI:** 10.64898/2026.02.11.705238

**Authors:** Ruidong Jia, Xinyi Zuo, Liting Zou, Li Wang, Letian Lin, Zhiqing Wang, Yuexia Zhang, Xingyu Chen, Fanrui Meng, Hantang Huang, Lan Lan, Zhimei Li, Fei Wang, Yuxuan Jin, Hongyan Shan, Rui Zhang, Hongzhi Kong, Peipei Wang

**Affiliations:** State Key Laboratory of Vegetable Biobreeding, Key Laboratory of Biology and Genetic Improvement of Flower Crops (North China), Ministry of Agriculture and Rural Affairs, Institute of Vegetables and Flowers, Chinese Academy of Agricultural Sciences, Beijing 100081, China; Kunpeng Institute of Modern Agriculture at Foshan, Foshan, Guangdong 528200, China; College of Horticulture, Northwest A&F University, Yangling, Shaanxi 712100, China; State Key Laboratory of Plant Diversity and Specialty Crops, Key Laboratory of Systematic and Evolutionary Botany, Institute of Botany, Chinese Academy of Sciences, Beijing 100093, China; China National Botanical Garden, Beijing 100093, China; University of Chinese Academy of Sciences, Beijing 100049, China; Guangdong Huabo Ecological Industry Co., Ltd., Foshan, Guangdong 528241, China; Bioinfo-Reach Bioinformatics Service, Guangzhou, Guangdong 510663, China; College of Plant Protection, South China Agricultural University, Guangzhou, Guangdong 510642, China; College of Horticulture, China Agricultural University, Beijing 100193, China; State Agricultural Biotechnology Centre (SABC), College of Science, Health, Engineering and Education, Murdoch University, Perth, Western Australia, 6150, Australia; Shanghai Jiaotong University, Shanghai, 200240, China

## Abstract

As an important ornamental crop, *Phalaenopsis* orchids exhibit extraordinary trait diversity with unclear genetic bases. Here, we present a haplotype-resolved, chromosome-level genome for an aneuploid cultivar ‘Santiago’. We reveal pervasive variation in the number and sequence among homoeologous genes. When serving as a reference, this genome facilitates the genetic understanding of trait variation in a hybrid population that is highly representative in trait polymorphisms. Specifically, nucleotide polymorphisms in a potential long-distance enhancer and the consequential expression variation of *PsAGL6-2*, presence/absence of *PsMYB12* and *b*-type homoeologous gene of *PsMYB2* determine lip morphology variation, presence/absence of venation-associated stripes and background pink color, respectively. Diverse functions of *PsMYBx1* in repressing anthocyanin accumulation across different cultivars further enhances the color patterning diversity in *Phalaenopsis*. Our study provides a practical framework for using a highly heterogeneous, haplotype-resolved genome to decode phenotypic diversity and has the potential to promote marker-assisted breeding in *Phalaenopsis*.

## Main

*Phalaenopsis* orchids are a major ornamental crop prized for their beauty and long flowering. Flowers in *Phalaenopsis* exhibit extraordinary diversity in organ organization, size, morphology, color patterning, scent, longevity, and so on, making *Phalaenopsis* a valuable system for horticultural research. Key genes regulating specific traits have been identified, such as floral scent^1,2^, floral organ identity^3–6^, color patterning^7–10^, petal size^11^, and cuticular wax^12^. However, the genetic bases of these diversity remain largely uncharacterized, except for the presence/absence (P/A) variation of floral scent^1,13^ and the variation in spot/patch patterns^7,9,10^. For example, the specific genetic variation responsible for the P/A of venation-associated stripes, diverse perianth colors and lip morphology across different cultivars are unclear.

To decipher the genetic bases of trait diversity, it is essential to associate allelic polymorphisms—including single nucleotide polymorphism (SNP), insertion and deletion (indel), copy number variation (CNV), and structural variant (SV)—with trait variation. A major bottleneck in *Phalaenopsis* is the lack of a chromosome-level reference genome; the existing genomes for *P. equestris* and *P. aphrodite* are only at the scaffold levels^14,15^. Consequently, the full scope of these allelic polymorphisms and their contributions to trait diversity is still poorly understood.

In this study, we present a haplotype-resolved, chromosome-level genome for a highly heterozygous aneuploid cultivar, *P.* ‘Santiago’ (hereafter *P.* sa, **Fig. 1A**). In addition, we generate a hybrid population (H80) which exhibits extensive and representative polymorphisms in lip morphology and floral color patterning (**Fig. S1**). By analyzing the whole genome resequencing data of this population using *P*. sa genome as the reference, we decipher genetic variation that account for these trait polymorphisms.

**Fig. 1.**
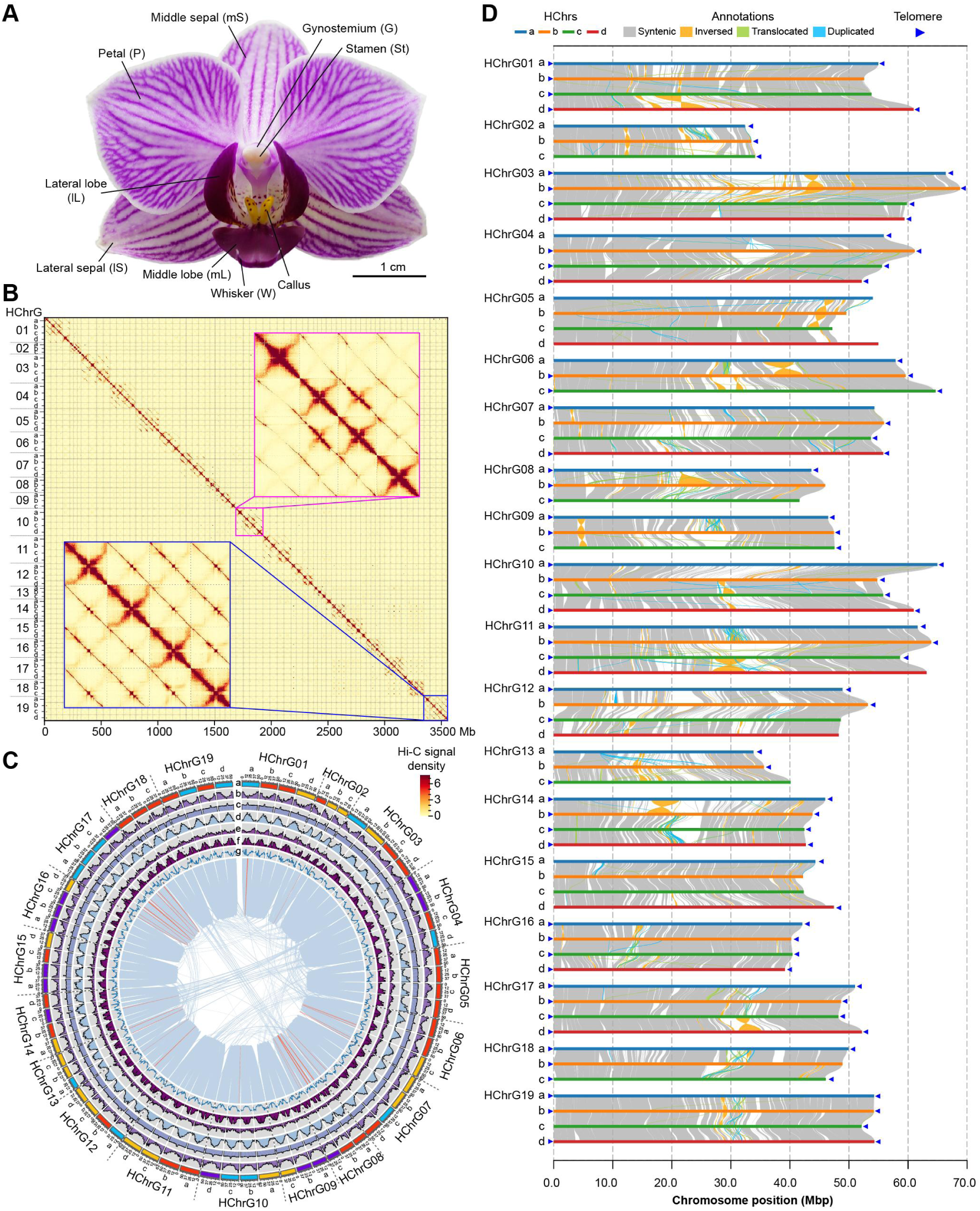
Genome assembly of *Phalaenopsis* ‘Santiago’ (*P.* sa). (**A**) A *P.* sa flower has one middle sepal, two lateral sepals, two petals, one lip, and a gynostemium that connects the stamen with two pollinia and the ovary. (**B**) Hi-C heatmap showing chromosomal interactions, with bin size of 250 Kb. Two insets illustrate weak interaction signals between homoeologous chromosomes (HChrs) of HChr group 10 (HChrG10, pink) and HChrG19 (blue) as examples. (**C**) Circos plot displaying genomic features within 1.5-Mb bins in *P.* sa. **a**, chromosome sizes, with different colors indicating predicted subgenomes, which is inferred from **Fig. S3**. **b**, gene density, ranging from 0 to 99. **c**, GC content, 15.38%–53.56%. **d**, transposable element coverage, 37.86%–99.79%. **e**, Copia LTR retrotransposon coverage, 0%–17.73%. **f**, Gypsy LTR retrotransposon coverage, 0%–98.51%. **g**, tandem repeat density, 0–251. The center region shows syntenic blocks between (blue lines) or within (orange lines) chromosomes. (**D**) Collinearity among homoeologous chromosomes, with predicted telomeres indicated.

## Results

### The genome assembly and annotation of *P.* sa

We generated the assembly of *P.* sa using a stepwise strategy (**Methods**). The whole genome size of *P.* sa was estimated to range from 3.3 Gb to 4.1 Gb by *k*-mer-based analyses and flow cytometry (**Methods**, **Fig. S2A,B**). *K*-mer frequency analysis (**Fig. S2C**) and karyotype analysis (70 chromosomes per cell, **Fig. S2D**) indicated *P*. sa as an aneuploid. Nineteen representative pseudo-chromosomes (a haplotype [1x] is supposed to contain 19 chromosomes)^15^ were first assembled using the PacBio HiFi reads and Hi-C short reads (**Methods**), to guide the following haplotype-resolved assembly. The primary unitigs were mapped to these 19 pseudo-chromosomes and grouped accordingly (**Methods**). Hi-C reads were used to scaffold these grouped unitigs to a haplotype-resolved, chromosome-level assembly per pseudo-chromosome. Finally, all these pseudo-chromosome-specific assemblies and unassigned unitigs were jointly refined and corrected using Hi-C reads (**Methods**).

The final assembly comprised 70 pseudo-chromosomes, with 6 and 13 homoeologous chromosome groups (HChrGs) having 3 and 4 haplotypes, respectively (**Fig. 1B,C**). The total assembly size was approximately 3.55 Gb, with a structural assembly quality indicator (AQI)^16^ of 96.57% and an assembly consensus quality value (QV)^17^ of 65.98. Benchmarking Universal Single-Copy Orthologs (BUSCO) assessment^18^ against the liliopsida database indicated 98.8% completeness, exceeding those of two other available *Phalaenopsis* genome assemblies (**Table S1**). Predicted telomere sequences were identified at most chromosome ends (**Fig. 1D**), further suggesting the relatively high assembly quality. Due to the complex hybrid history, the *P.* sa genome did not display obvious phases of subgenomes (**Fig. 1C, Fig. S3**). We predicted 93,708 protein-coding genes using *de novo*, homology-based, and transcriptome-supported approaches (**Methods**, **Table S2**), with a BUSCO (liliopsida) completeness of 98.5%. Repetitive sequences constituted 67.97% (2.41 Gb) of *P. sa* genome (**Table S3**). Additionally, the genome annotation included 34,366 non-coding RNAs (**Table S4**).

### Homoeologous gene variation in *P.* sa

Syntenic analysis (**Methods**) revealed extensive structural variation between HChrs (**Fig. 1D**, **Table S5**). Accordingly, all the 93,708 protein-coding genes were classified into 33,697 homoeologous gene groups (HGGs) (**Methods**), which were further categorized into six types according to variation in the number and sequence among HGs (**Fig. 2A**). Genes in type 1 contained a single HG each; type 2 genes had ≥2 HGs while with HChr-specific HG losses; type 3 genes had HChr-specific tandem duplicates; type 4 genes had the same number of tandem duplicates per HChr; type 5 genes had single copy per HChr but with different protein domain constitutions; type 6 genes had single copy per HChr with same domain constitutions (**Fig. 2A**). Protein domain constitutions (**Methods**) were examined because proteins with different domain constitutions are likely to perform divergent functions^19,20^.

**Fig. 2.**
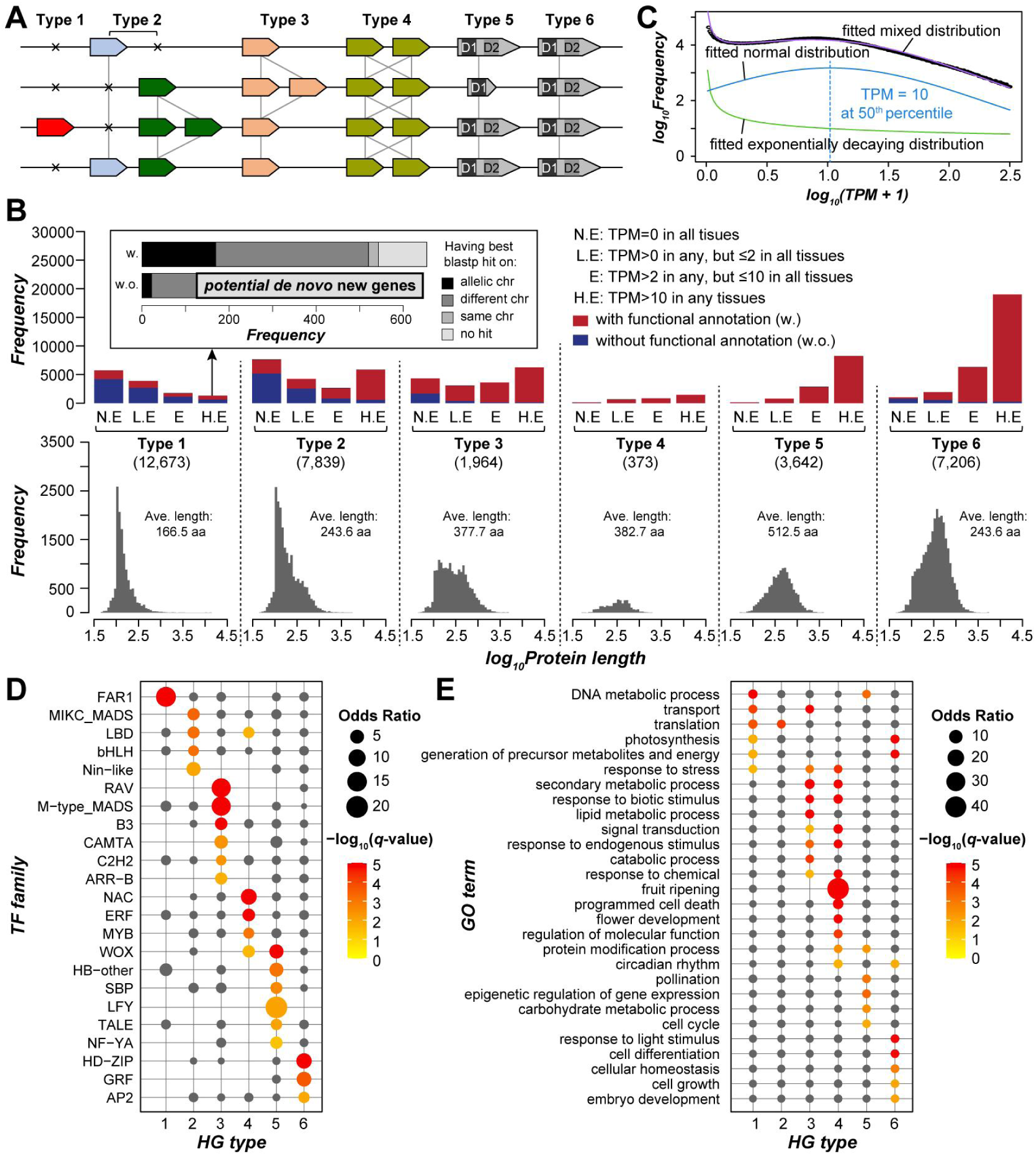
Homoeologous gene variation in *P.* sa genome. (**A**) Diagram illustrating the six HG types. D1 and D2: protein domain 1 and domain 2. (**B**) Frequency of six HG types (upper panel) with the number of HGGs listed within the parentheses, and their protein length distribution (lower) with the average length per type indicated as well. Box colors indicate genes with or without functional annotations. TPM: Transcripts Per Kilobase; N.E.: not expressed; L.E.: lowly expressed; E.: expressed; H.E.: highly expressed. **Inset**: frequency of highly expressed (TPM>10) type 1 genes with (w.) or without (w.o.) functional annotation. Different gray scales indicate whether a gene has the best non-self blastp hit on different, homoeologous or same chromosomes, or has no blastp hit. (**C**) Mixed distribution of gene expression levels, illustrating how TPM=10 was selected as the cutoff threshold for high expression (see **Methods**). Fitted normal distribution indicates true expression, while fitted exponentially decaying distribution indicates noise expression. (**D,E**) Enriched transcription factors (**D**) and biological_process GO terms (**E**) in six HG types based on Fisher’s exact test. Dot size: odds ratio; dot color: -log_10_ *q*-values which are the adjusted *p*-values using the “BH” method; cases with *q*-values≥0.05 or odds ratio<1 were colored gray.

Genes in types 1-6 differed in terms of expression pattern, functional annotation (**Methods**) and encoded protein length (**Fig. 2B,C**). They tended to belong to different transcription factor (TF) families (**Fig. 2D**) and perform different biological functions (**Fig. 2E**). Specifically, type 1 genes tended to be not or lowly expressed in all 28 examined samples (**Table S6**), have no functional annotations and have relatively short proteins (**Fig. 2B**). All the 13 *FAR-RED IMPAIRED RESPONSE 1 (FAR1*) genes in type 1 (**Fig. 2D**) were not or lowly expressed (**Fig. S4**), indicating the ultimate destiny of being lost from the genome. *FAR1* genes are essential for phytochrome signaling and balancing plant growth and defense responses under shade conditions^21^. The deletion of certain *FAR1* genes from the *Phalaenopsis* genome may be results of adaptation to shady, damp forest understory. In contrast, some other type 1 genes might be *de novo* new genes endowing *Phalaenopsis* with new phenotypic and adaptive characteristics: 519 genes were highly expressed (TPM>10) in certain tissues while having no functional annotations and no non-self blastp hits within *P.* sa genome (**Fig. 2B**). Among them, 243 (46.8%) were preferentially expressed in stamen (**Fig. S5A-C**), consistent with previous findings that new genes tended to arise from male reproductive organs^22,23^. The remaining 276 genes were preferentially expressed in other tissues (**Fig. S5D-G**), suggestive of their potential functions in the corresponding tissues.

Type 2 genes were enriched for TFs that play important roles in floral development, cell proliferation, determination and differentiation, lateral organ formation, responses to environmental stimuli, and nitrate responses (**Fig. 2D**). Type 3 genes were enriched for TFs and GO terms involved in flowering, embryogenesis, stress responses and signaling (**Fig. 2D,E**). Type 4 genes were enriched in TFs and functions related to plant growth and stress responses (**Fig. 2D,E**). Genes in types 5 and 6 were enriched for TFs responsible for and GO terms related to essential and conserved functions (**Fig. 2D,E**). These results indicate that the HChr-specific gene duplicates/losses (types 2,3), together with those tandem duplicates retained in all HChrs (type 4), may have driven the high diversity of floral traits and the adaptation of *Phalaenopsis* to diverse environments.

### Dosage effects of homoeologous gene count

Considering the aneuploidy nature of and the substantial HG count variation in *P*. sa genome, we asked whether the chromosome ploidy (3x vs 4x) and HG count variation affect the total expression amount per gene. To ensure a valid comparison, only reciprocal best-matched (RBM) gene pairs with different HG counts—each gene had no HChr-specific tandem duplicates and had the same protein domain constitutions among HGs—were examined (C1-C8, **Fig. 3A**). This strategy was adopted because the gene expression in the hybrid parents of *P.* sa were not available. Gene expression scaled with chromosome ploidy: summed expression levels of type 6 genes with 4 HGs (4/4) were generally higher than those of their 3/3 RBM counterparts (**Fig. 3B**), but became comparable when averaged per chromosome count (**Fig. 3C**). This dosage effect was similar to those found in other aneuploid organisms, where gene expression in aneuploids were compared with their corresponding diploids, such as maize^24^, budding yeast^25^ and *Candida albicans*^26^.

**Fig. 3.**
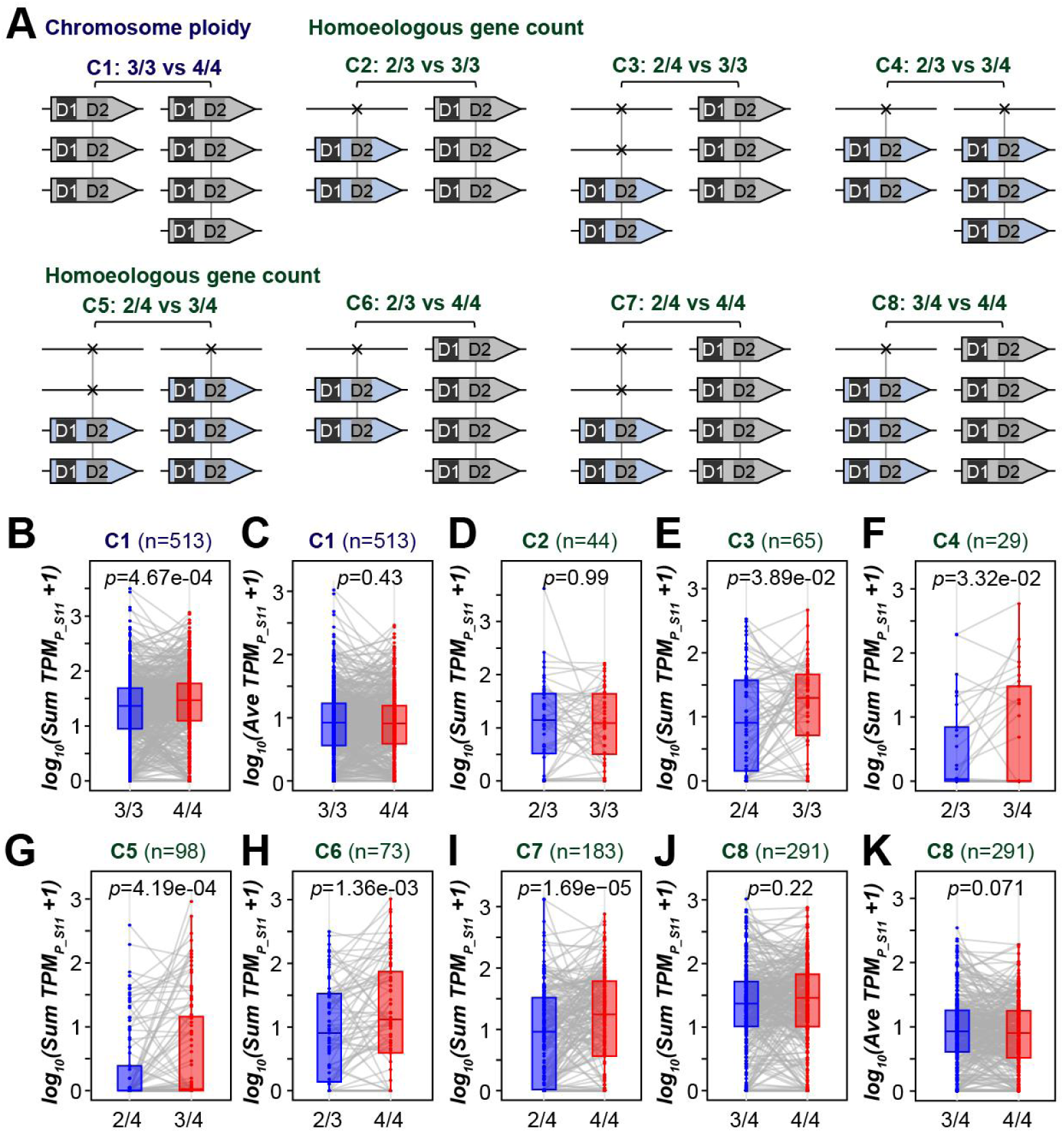
Dosage effects of chromosome ploidy and homoeologous gene count on gene expression. (**A**) Diagrams showing eight comparisons, where C1 compares reciprocal best-matched (RBM) gene pairs located on HChrGs with different ploidies (3x vs 4x), while C2-C8 compare RBM genes with different HG counts. 2/3 and 2/4: only two HGs were retained for HChrGs with 3x and 4x, respectively; 3/4: three HGs were retained for HChrGs with 4x. (**B,C**) Summed (**B**) and averaged (**C**) HG expression levels in petals at S11 of RBM gene pairs between HChrGs with 3x or 4x. (**D-J**) Summed HG expression levels for RBM gene pairs in C2-C8. (**K**) Averaged HG expression levels for RBM gene pairs in C8. Regarding (**B-K**), see **Dataset 1** for expression results in other tissues.

In addition to chromosome ploidy, HG counts also had dosage effects on gene expression as demonstrated by all but two comparisons (C3-C7, **Fig. 3E-I**): genes with more HGs tended to have higher expression levels than their RBM counterparts with fewer HGs. The other two comparisons (C2, C8) indicated expression compensation when only one HG was lost: no significant differences in expression between 2/3 and 3/3 genes, and between 3/4 and 4/4 genes (**Fig. 3D,J)**, but significantly higher averaged levels of 3/4 genes than 4/4 genes (*p*=0.071, corresponding to *p*=0.035 of one-sided Wilcoxon rank sum test, **Fig. 3K**). These results suggest that buffering mechanisms counteract the disruption of homeostasis caused by immediate HChr-specific gene losses (i.e., 2/3 and 3/4 genes), but not that caused by chromosome inequivalence.

### Genetic bases of key trait polymorphisms

The extensive HG variation in *P*. sa were mainly derived from allelic polymorphisms of its hybrid parents, which represented part of the entire polymorphism set of *Phalaenopsis* that contribute to the trait diversity. To evaluate the utility of *P*. sa genome in revealing the genetic bases of those diversity, we generated a hybrid population (H80) that displayed substantial variation in key floral traits (**Fig. 4A, Fig. S1**), especially in lip morphology and color patterning. Different trait categories can combine randomly, resulting in a unique phenotype for each individual in this population. This makes it a highly representative set of *Phalaenopsis* orchids. We resequenced the genomes of 46 H80 representative individuals and performed genome-wide association study (GWAS) using *P*. sa genome as the reference.

**Fig. 4.**
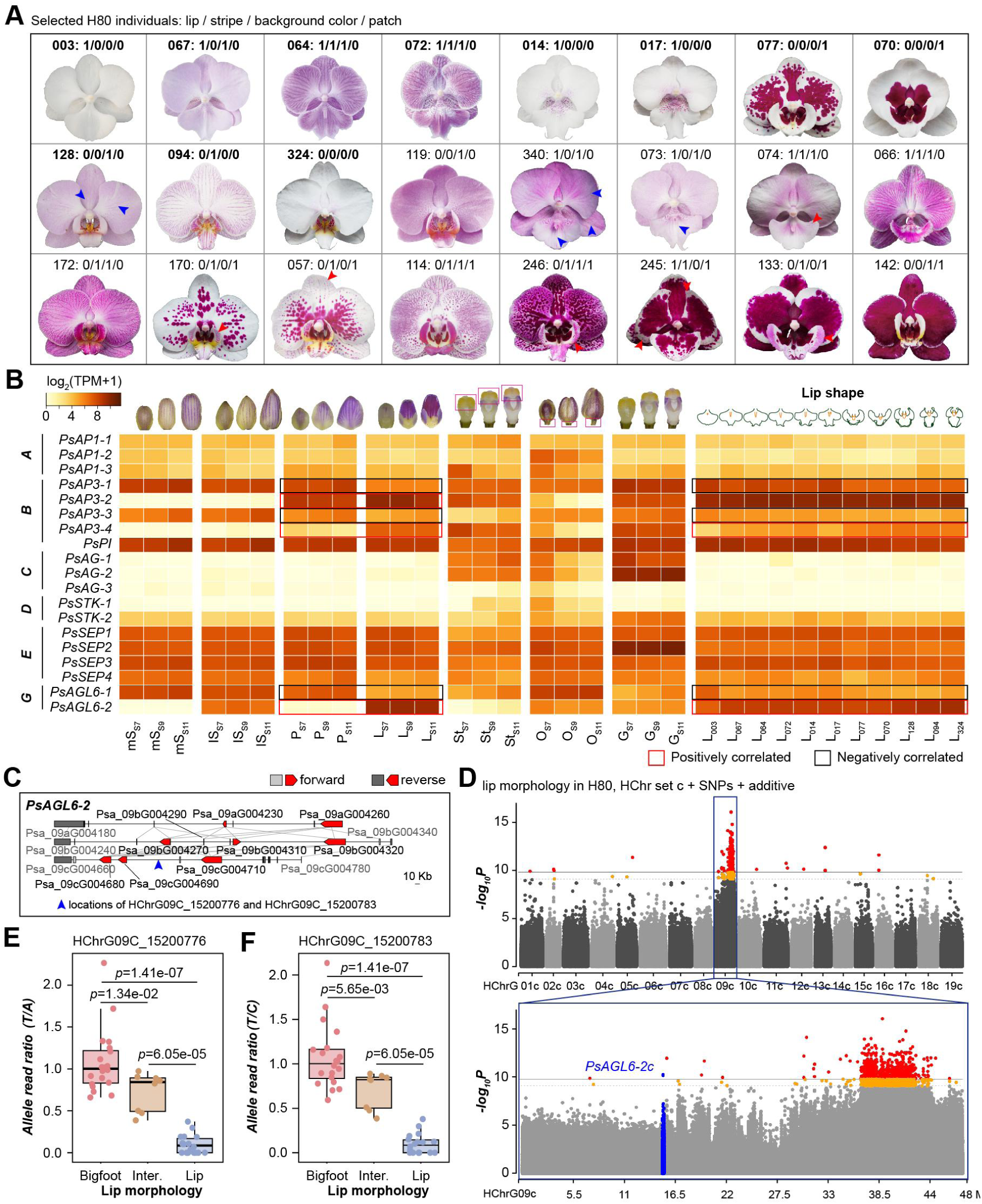
Floral trait variation in H80 and the genetic basis of lip morphology variation. (**A**) Phenotypes of 24 H80 individuals (see **Fig. S1** for all 46 individuals). The category of each individual in lip morphology (0: lip, 1: petal-like organ), stripe P/A (0: A, 1: P), background color (0: white, 1: pink), and spot/patch types (0: spotted or non-spotted, 1: patched) were indicated. Bold font: individuals with transcriptome data of the lip. Red arrowhead indicates stripe, while blue arrowhead indicates stripe-like shadow. (**B**) Heatmap showing expression profile of A-G classes of MIKC-type MADS-box genes in different floral organs at three stages (S7, S9 and S11), and in the S9 lips of 11 representative H80 individuals. Color scale indicates log-transformed expression levels. mS: middle sepals; lS: lateral sepals; P: petals; L: lip; St: stamen; O: ovary; G: gynostemium. Red and black boxes: expression of genes positively or negatively correlated with petal-to-lip transitions, respectively. (**C**) HG variation of *PsAGL6-2*. Red boxes: *PsAGL6-2* genes with directions indicated; light or dark gray boxes: neighboring genes with forward or reverse directions, respectively; boxes linked using gray lines between HChrs: HGs. The blue arrowhead indicates locations of two significant SNPs. (**D**) Manhattan plot for GWAS of lip morphology in H80, using SNPs referred to HChr set c with the additive marker-effect model. Horizontal solid and dashed lines indicate -log_10_ (0.01/variant counts) and -log_10_ (0.05/variant counts) (**Methods**), respectively; red and orange dots denote SNPs significantly associated with lip morphology variation; blue dots represent SNPs located within or adjacent (±2 kb) to the *PsAGL6-2* gene clusters. (**E**,**F**) Allele read ratio of two SNPs (count of reads supporting “T” to count of reads supporting “A” in **E**, and count of reads supporting “T” to that of “C” in **F**) within H80 individuals with different lip morphologies. *P*-values are from two-sided Wilcoxon rank-sum tests.

Key regulators of these traits have been previously identified (see **Table S7** for their gene IDs in the *P.* sa genome), including *PeAGL6-2* (*AGAMOUS-LIKE 6-2*) in lip organ identity determination^27^, *PeMYB2*, *PeMYB12* and *PeMYB11* in the formation of floral background color, venation-associated stripes and spots, respectively^7^, and *MYBx1* in repressing anthocyanin accumulation in non-patched regions^8–10^. Their functions were relatively conserved in *P*. sa and H80 according to their expression profiles (**Fig. 4B**, **Fig. 5A,B**). Thus, we focused on identifying genetic variants within or near these genes that were significantly associated with corresponding trait polymorphisms, and assessing how they contributed to trait differentiation in H80. Since H80 individuals were shown to be aneuploids as well by karyotype analysis (**Fig. S2F**, 2n=64, >3x), they were treated as tetraploids in the GWAS to facilitate the analysis. Moreover, given that 70 chromosomes in *P.* sa can not be readily phased into subgenomes (**Fig. 1C**), the original HChr designations (such as that HChrG01a∼HChrG19a were designated as HChr set a) were retained when serving as the reference genome during the GWAS.

**Fig. 5.**
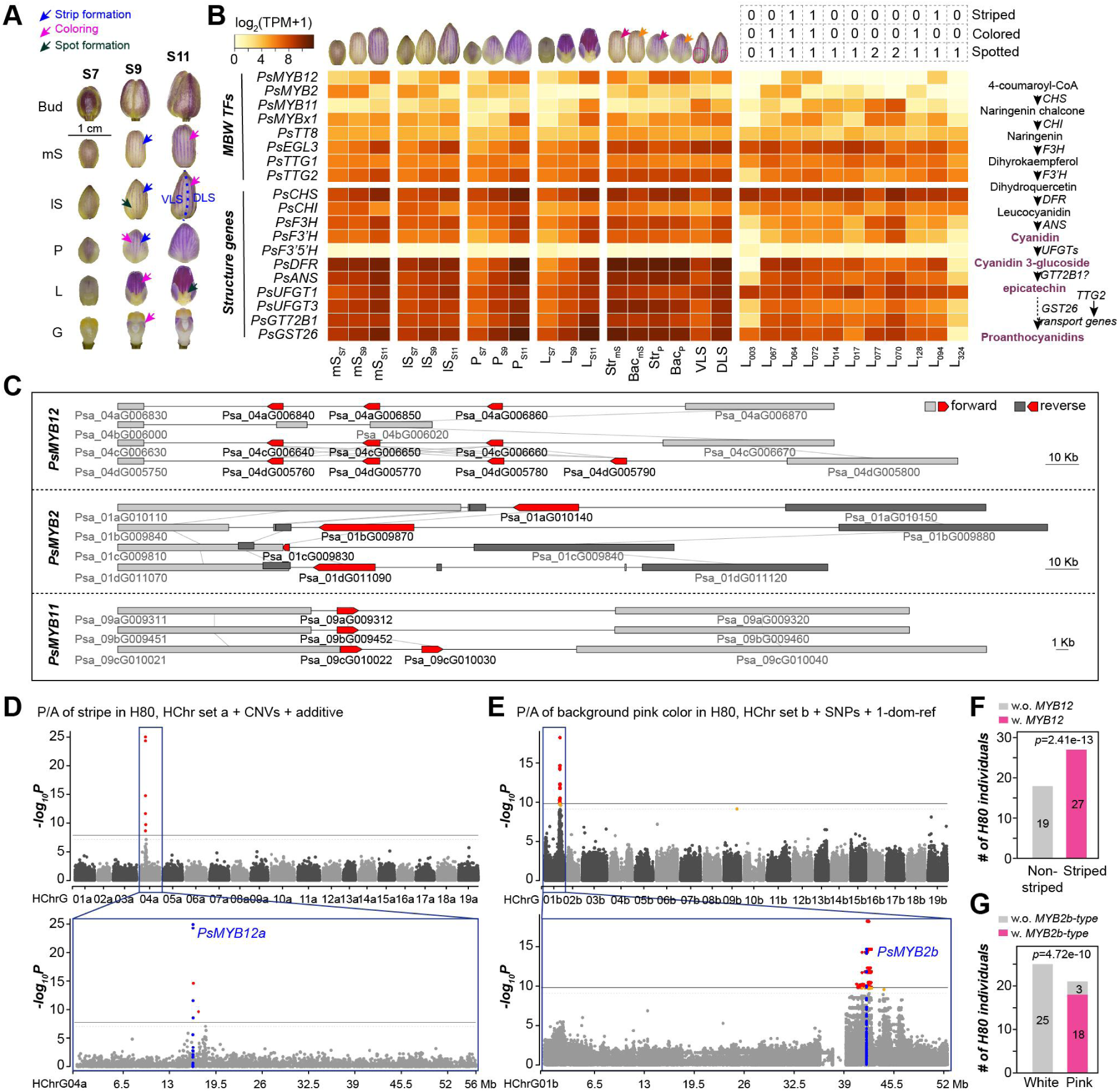
Genetic bases of color patterning variation in H80. (**A**) Floral organs at three developmental stages showing important events of color patterning. VLS and DLS: ventral and dorsal half of the lateral sepals, respectively. (**B**) Heatmap showing expression profiles of genes participating in color patterning. The classifications of each H80 individual in the presence (0) or absence (1) of stripes, background pink colors and spots (2 indicates patches) are listed. MBW TFs: transcription factors that form the MYB-bHLH-WD40 complexes. The biosynthesis pathway for the prominent anthocyanin in *Phalaenopsis* orchids—cyanidin—is shown on the right. Str: stripe; Bac: inter-stripe background region. (**C**) HG variation of *PsMYB12*, *PsMYB2* and *PsMYB11*. (**D**) Manhattan plot for GWAS of stripe P/A in H80, using CNVs referred to HChr set a with the additive marker-effect model. (**E**) Manhattan plot for GWAS of background color in H80, using SNPs referred to HChr set b with the 1-dom-ref marker-effect model. (**F,G**) Association of P/A of *PsMYB12* genes (**F**) and P/A of *PsMYB2b*-type sequences (**G**) with stripe P/A and background color, respectively. *P*-values are from Fisher’s exact test.

### Lip morphology variation and nucleotide polymorphisms nearby *PsAGL6-2* genes

*PsAGL6-2* had 2-4 tandemly arranged duplicates on three HChrs (corresponding to HGG type 3, **Fig. 4C**), with only one copy encoding a full-length protein per HChr (**Fig. S6**). GWAS analysis for the lip morphology (lips vs petal-like organs) identified two significant SNPs in the intergenic region between two *PsAGL6-2* copies (**Fig. 4C**) only when HChr set c was used as the reference (**Fig. 4D, Fig. S7-9**). The allele read ratios (the number of reads supporting one HG divided by the number of reads supporting the other HG) of these two SNPs were significantly associated with lip morphology in H80 (**Fig. 4E,F**, **Table S8**). Expression levels of *PsAGL6-2* showed positive correlation with lip morphology (**Fig. 4B**). These results suggest that the transformation of lips into petal-like organs in H80 is associated with reduced expression of *PsAGL6-2*, potentially due to the introduction of an allele containing mutations in a long-distance enhancer (∼65 Kb upstream of *Psa_09cG004690* and ∼86 Kb downstream of *Psa_09cG004710*, **Fig. 4C**). How these mutations reduced *PsAGL6-2* expression remained to be determined, potentially through genome editing of these loci in cultivars bearing lips or petal-like organs. Additionally, expression of other proposed lip (*PsAPETALA3-4* [*PsAP3-4*]) and petal identity genes (*PsAP3-1*, *PsAP3-3*, and *PsAGL6-1*; see **Fig. S10** for the gene tree of MADS-box genes)^28^ exhibited positive and negative correlations with petal-to-lip transitions, respectively (**Fig. 4B**). These results suggest that expression of these genes might be regulated by *PsAGL6-2*, thereby coordinating lip morphogenesis.

### Presence/absence of stripes and presence/absence of *PsMYB12* genes

Expression of *PsMYB12* genes was highly associated with stripe formation in *P.* sa and the presence of stripes in H80 population (**Fig. 5A,B**). It lost its entire tandemly duplicated gene cluster from HChrG04b (HGG type 2, **Fig. 5C**). CNV-based GWAS for P/A of stripes (**Fig. 5D, Methods**) using HChr set a as the reference identified significant variants located in and nearby *PsMYB12* genes while with the fewest false positives compared with others (**Fig. S11**). Consistent with this, stripes P/A in H80 was perfectly associated with P/A of resequencing reads mapped to *PsMYB12* genes (**Fig. 5F**, **Fig. S12, Table S9**), suggesting that stripe P/A were determined by *PeMYB12* P/A.

Since we didn’t identify any genes orthologous to *PsMYB12* in the other eight orchid species (including the non-striped *P. aphrodite*) except for *P. equestris* (**Fig. S13A**), we asked when and how the *PsMYB12* genes and stripes originated in *Phalaenopsis*. Interestingly, stripes were only observed for the branch leading to *P. equestris* and *P. lindenii* in wild *Phalaenopsis* species (**Fig. S13B**). To test if *PsMYB12* genes originated before or after the most recent common ancestor (MRCA) of *Phalaenopsis*, we resequenced the *Vanda* ‘Pakchong Blue’, a stripe-bearing orchid cultivar outside *Phalaenopsis*. We detected reads mapped to *PsMYB12* genes, albeit with a high degree of divergence within and near these genes (**Fig. S13C**). Given that other orchids like *Paphiopedilum* also have stripes, these results suggest that *MYB12* genes originated in other orchids but were absent in the MRCA of *Phalaenopsis* and were introduced into the MRCA of *P. equestris* and *P. lindenii*, potentially via gene introgression.

### Background pink color and the *b*-type HG of *PsMYB2*

*PsMYB2* had a single copy per HChr (**Fig. 5C**), but the one on HChrG01c (*Psa_01cG009830*) had lost its R2 domain (HGG type 5, **Fig. S14A**). GWAS for the P/A of background pink color using 1-dom-ref model (**Methods**) with HChr set b as the reference identified significant SNPs and indels located in and nearby *PsMYB2* genes while with the fewest false positives among others (**Fig. 5E**, **Fig. S15-17**). In this 1-dom-ref model, the *PsMYB2b*-type HG was considered as dominant in determining the pink color, consistent with its higher expression levels than other HGs (**Fig. S14B**). The P/A of *PsMYB2b*-type sequences can nearly perfectly predict the P/A of background pink color on sepals/petals for all but three individuals—H80-066, H80-072 and H80-114 (**Fig. 5G**, **Table S10**). The background color in these exceptions may be governed by other non-*PsMYB2* genes, since *PsMYB2* expression was almost undetectable in the lip of H80-072 (the summed TPM of four HGs was 1.32, **Fig. 5B**, **Fig. S14B**). The protein sequence of PsMYB2b differed from PsMYB2a by only two amino acids in the C-terminal region (**Fig. S14A**), suggesting that the pink color in *P.* sa and most H80 individuals with pink flowers was likely determined by the *PsMYB2b*-specific expression. It requires further investigation of which region of the *PsMYB2b* promoter drives its specific expression.

### Spot/patch pattern variation

*PsMYB11* had an extra tandem copy (*Psa_09cG010030*) on HChrG09c (HGG type 3, **Fig. 5C**), which harbored a few mutations in protein sequences and the upstream sequences (**Fig. S14C,D**) compared with other HGs. Expression of *PsMYB11* genes, especially *Psa_09cG010030*, was highly associated with the spot formation in *P*. sa and the spot/patch pattern variation in H80 (**Fig. 4A**, **Fig. S14E**). GWAS for the P/A of patches identified numerous variants distributed across the entire genome, but no variants were within or near the *PsMYB11* genes (>96.8 Kb), regardless of variant types used and HChr sets referred to (**Fig. S18-20**). Given that the patch formation was reported to be determined by the *HORT1* insertion in the promoter sequences of *PeMYB11* genes^9^, and that *P.* sa flowers harbored only small spots, the failure to identify *PsMYB11*-related variants is potentially due to the lack of *HORT1* insertion in *PsMYB11* promoters.

### Diverse functions of *PsMYBx1* in repressing anthocyanin accumulation

*PsMYBx1* had only one HG (HGG type 1, **Fig. 6A**). Its expression was associated with both stripe and spot/patch formation in *P.* sa and H80 (**Fig. 5A,B**), indicating potentially dual function of *PsMYBx1* in repressing anthocyanin accumulation in both inter-stripe and inter-spot/patch regions. This dual function was supported by the disruption of stripes in regions surrounding patches, such as in H80-057, H80-114, and H80-245 (**Fig. 4A**). Given this complexity of color patterning imposed by *PsMYBx1* genes, no single color patterning variation can be readily determined to be related to *PsMYBx1*. Thus, instead of performing GWAS, we silenced *MYBx1* genes via virus-induced gene silencing (**Methods**) in another cultivar, *P*. ‘Jinbianlinglong’ (*P.* ji) (**Fig. 6B,C**), for which substantial seedlings are available and the flowering regulation is relatively readily. *MYBx1* had four HGs in *P*. ji (**Fig. S21B**), compared with a single HG in *P*. sa. *PjPDS* (*Phytoene desaturase*) was co-silenced to indicate regions with gene silencing phenotypes where the yellow-green color was faded (**Fig. 6B**).

**Fig. 6.**
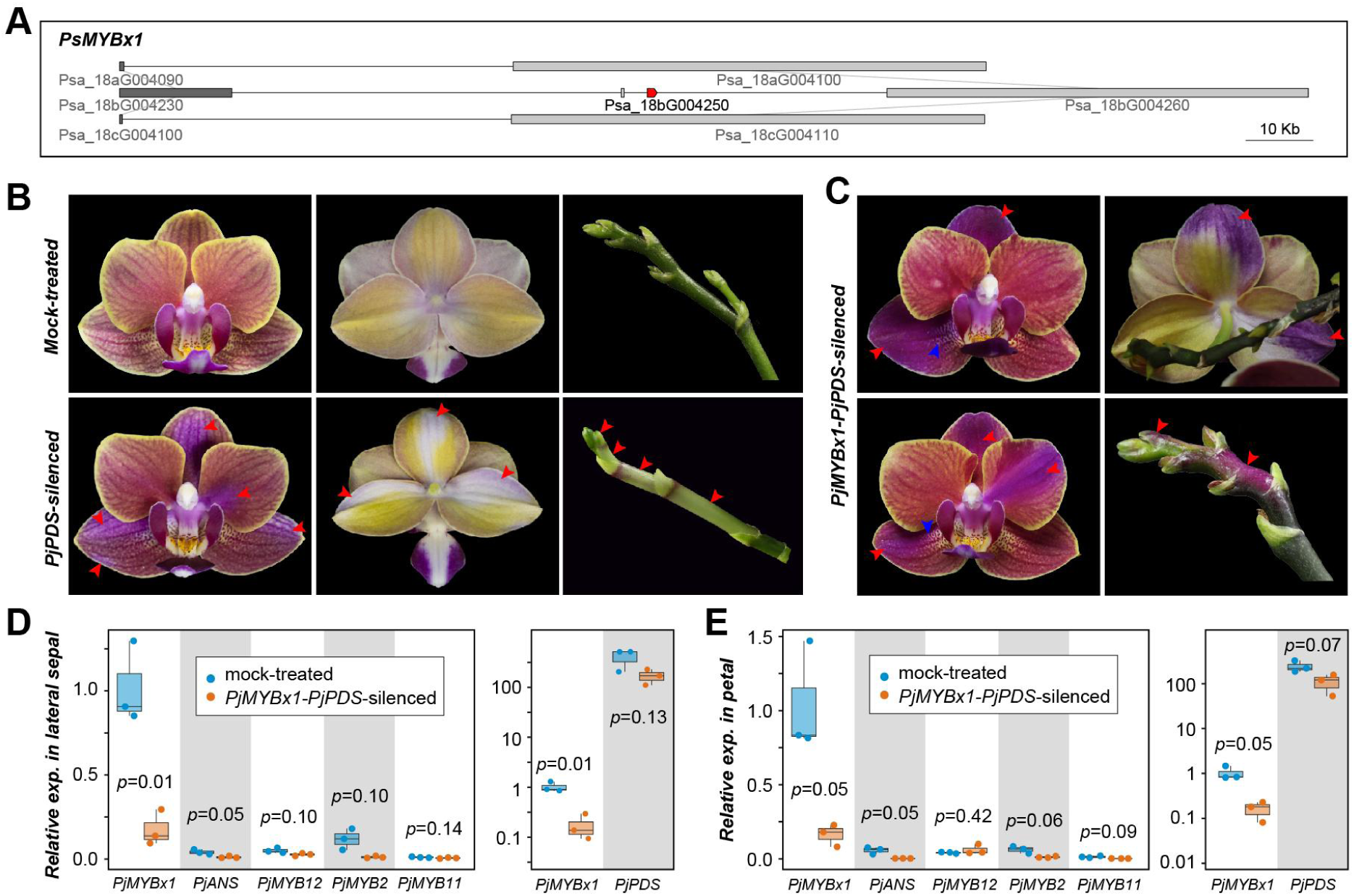
Functions of *PjMYBx1* in repressing anthocyanin accumulation in diverse regions. (**A**) HG variation of *PsMYBx1*. (**B**) Flower and inflorescence images showing phenotypes of mock-treated (upper panel) and *PjPDS*-silenced (lower panel) plants. (**C**) Flower and inflorescence images of *PjMYBx1*-*PjPDS*-silenced plants. Red arrowheads indicate regions with phenotypic changes when compared with mock-treated plants, while the blue ones indicate the unchanged inter-spot regions. (**D,E**) Expression of color patterning-related genes in mock-treated and *PjMYBx1*-*PjPDS*-silenced sepals (**D**) and petals (**E**). *P*-values are from t-test.

In *PjMYBx1*-*PjPDS*-silenced plants, ectopic anthocyanin pigmentation was observed on the inflorescence stalk, abaxial surface of perianth, and in the inter-stripe regions (**Fig. 6C**). However, the inter-spot regions remained unchanged (blue arrowheads in **Fig. 6C** where the yellow-green color was faded), contrasting with the proposed repressing role of *MYBx1* in these regions^10^. Although expression of color patterning-related genes examined were generally decreased in *PjMYBxl*-*PjPDS*-silenced sepals (**Fig. 6D**) and petals (**Fig. 6E**) compared with those in mock-treated plants, a slight increase in *PjMYB12* expression was detected in petals. Taken together, these results demonstrate the multiple functions of *MYBx1* in repressing anthocyanin accumulation in diverse regions, and suggest variation in its functions across different *Phalaenopsis* orchids.

## Discussion

The haplotype-resolved, chromosome-level genome of *P*. sa served as a suitable reference in revealing the genetic bases of trait diversity in *Phalaenopsis*. Given the high heterogeneity of *P*. sa genome and the diverse genetic mechanisms underlying different trait variation, we demonstrated that the choice of HChr sets as the reference genome, variant types, and marker-effect models is critical for revealing the genetic bases of target trait variation. The ratio of true to false positives of the identified significant variants can be the primary metric for optimization. Specifically, for lip morphology variation, stripe P/A variation, and pink color P/A variation, the optimal models were HChr set c + SNPs + additive model, HChr set a + CNVs + additive model, HChr set b + SNPs/indels + 1-dom-ref model, respectively. However, for spot/patch variation, genomes of patch-bearing *Phalaenopsis* cultivars are supposed to serve as superior references compared with *P*. sa genome.

Although genes regulating some of the key traits in *Phalaenopsis* have been well established^1,2,7,12,27,29,30^, what and how allelic polymorphisms of these genes have contributed to the extraordinary trait diversity remain underexplored, restricting their utilization in breeding practices. The genetic bases of three key trait variation revealed in this study—nucleotide polymorphisms nearby *PsAGL6-2* genes, P/A of *PsMYB12* genes, and P/A of *b*-type HG of *PsMYB2*—enable design of a multiplex polymerase chain reaction (MPCR) kit for early-stage prediction of these three traits simultaneously. Given the unusually long juvenile phases of *Phalaenopsis* orchids (3-5 years), such a MPCR kit has great potential in substantially shortening breeding cycles and reducing breeding costs.

## Methods

### Flow cytometry analysis

Young leaves of *P.* sa (ANTHURA BV, with US patent US20150047084P1) seedlings were used for the flow cytometry analysis, with young leaves of cultivated tomato one month after seed germination being used as the internal reference. Suspensions of *P.* sa and tomato leaves were strained using propidium iodide and mixed, and a BD FACScalibur flow cytometer was used to detect the stained cell nuclei.

### Karyotype analysis

Root tips of *P.* sa were harvested between 9:00 and 10:00 a.m., and approximately 0.5 cm segments were excised using a clean razor blade. These segments were incubated in 0.1% colchicine solution at room temperature in the dark for 5 h to induce chromosome condensation, and were then fixed in Carnoy’s solution (absolute ethanol:glacial acetic acid = 3:1, v/v) overnight. Fixed samples were rinsed three times with distilled water, dehydrated in 75%, 85%, 95%, and 100% ethanol for 10 minutes each, and then were stored in 75% ethanol at 4 °C. To facilitate cell wall separation, pretreated samples were incubated in 75 mM KCl solution at 37 °C for 30–60 minutes, and treated with 1 M HCl at 60 °C in a water bath for 10 minutes. Samples were stained with carbol fuchsin solution (Coolaber, Beijing, China) for 20 minutes, squashed on microscope slides, observed and photographed using a 60× objective lens on an OLYMPUS BX53 optical microscope.

### Genome sequencing and Hi-C sequencing

For HiFi sequences, the total DNA was extracted from *P.* sa young leaves, and used for the library construction and genome sequencing with the PacBio Sequel II/Sequel IIe platform, following the manufacturer’s instructions at Haorui Genomics (Xi’an, China). Approximately 72.1 Gb of PacBio HiFi reads were generated using the CCS mode.

For Hi-C sequences, the frozen young leaves were ground with liquid nitrogen and then fixed with 1% formaldehyde in vacuum at room temperature for 0.5-1h to preserve the three-dimensional chromatin architecture, and 0.15 M glycine was added in the solution for 5 min to quench the reaction. The mixed solution was then filtered with a 40 nm cell filter.

Chromatin was digested with 200 U DPNII restriction enzyme (Qiagen, 59971) at 37 ℃ for 16 h. DNA ends were labeled with biotin and incubated at 37 ℃ for 45 min. DNA ligation was performed by the addition of T4 DNA ligase and incubation for 1-2 h. Then proteinase K was added to reverse cross-linking during incubation at 65 ℃ for 1-2 h. DNA fragments were purified and dissolved in 86 μL of water. Purified DNA was fragmented to a size of 350-500 bp, and DNA ends were then repaired. DNA fragments labeled with biotin were finally separated on Dynabeads® M-280 Streptavidin (Life Technologies, 60210). The sequencing was performed with the Illumina NovaSeq 6000 for paired-end sequences with 150 bp read length. A total of ∼130.6 Gb raw data was produced. After filtering adapter sequences and read pairs with > 10% N or with low quality (Q ≤ 5) for >50% bases in at least one read, ∼123.3 Gb clean data was retained.

### Genome survey

The genome size was estimated via *k*-mer frequency analysis using the PacBio HiFi reads. The Jellyfish software (v2.2.10)^31^ was used to generate the counts of 31-mers, and then the GenomeScope2 software (v2.0)^32^ and Genomic Character Estimator (GCE) software (v1.0.2)^33^ were applied to estimate the genome size and heterozygosity. SmudgePlot software (v0.2.2)^32^ was used to estimate the ploidy level of *P.* sa genome.

### *De novo* genome assembly

Since it lacks a chromosome-level reference assembly for *Phalaenopsis* species, at the first place, we attempted to assemble a set of draft representative pseudo-chromosomes to guide the subsequent refined assembling. Three strategies were conducted to assemble the draft genome: (**ⅰ**) all the PacBio HiFi reads were assembled using HiFiasm program (v0.19.6-r597)^34^ with the parameter ‘l=0’ (to disable the purge of duplication); (**ⅱ**) the PacBio HiFi reads were assembled using HiFiasm with the parameters ‘l=3, hom-cov=57, hg-size=1.3g, n-hap=3, s=0.3, D=8, n-weight=6, n-perturb=80,000, f-perturb=0.3’, with the assumption that the *P.* sa was a triploid, and the Hi-C reads were integrated to achieve haplotype-resolved assemblies; (**ⅲ**) the PacBio HiFi reads were assembled with parameters ‘l=3, hom-cov=39, hg-size=1.5g, n-hap=4, s=0.3, D=8, n-weight=6, n-perturb=80,000, f-perturb=0.3’, with the assumption that the *P.* sa was a tetraploid, and the Hi-C reads were also integrated.

The Hi-C reads were mapped to the resulting primary unitigs from assembling **ⅰ**, and haplotigs of each haplotype from assembling **ⅱ** and **ⅲ**, separately, using bwa-mem2 (v2.2.1)^35^. The resulting bam files were further filtered using the ‘filter_bam.py’ utility in the HapHic (v1.0.2)^36^ with the parameters ‘mapq=1, NM=3’. The filtered bam files were then fed into the HapHic pipeline with parameters ‘correct_nrounds=2, quick_view=T’, and the output files .hic and .assembly were imported into the Juicebox Assembly Tools (JBAT, v1.08)^37^. Primary unitigs from strategy (i) and haplotigs from (ii) and (iii) were then ordered, joined, and manually adjusted based on Hi-C interaction patterns to obtain scaffolds, with the prior knowledge that nearby regions within the same chromosome having strong Hi-C signals whereas allelic regions having subdominant signals. The resulting scaffolds from all three strategies were compared, and 19 were selected as representative pseudo-chromosomes based on Hi-C signal continuity and Benchmarking Universal Single-Copy Orthologs (BUSCO) ^18^ completeness. These 19 pseudo-chromosomes—seven from assembly **i**, ten from **ii**, and two from **iii**—constitute the draft reference assembly (assembly v1.0).

Next, using the v1 draft assembly as a reference, we mapped all raw unitigs from assembly **ii** and grouped them into 19 chromosomal groups. Unitigs within each group were ordered, joined, and manually adjusted based on Hi-C interaction patterns, with the parameters ‘mapq=0, NM=3’ for filter_bam.py. This produced 70 preliminary pseudo-chromosomes. Remaining unassigned unitigs were then incorporated and the entire set was reviewed and corrected using Hi-C signals to yield the final assembly (assembly v2.0). SubPhase (v1.2.6)^38^ was used to phase subgenomes based on potential subgenome-specific *k*-mers, with parameters “-k 15 -q 50 -f 12”.

### Genome assembly quality assessment

The *P.* sa genome assembly contiguity was accessed using Quality Assessment Tool for Genome Assemblies (QUAST, v5.0.2)^39^. The BUSCO completeness was conducted using BUSCO (v5.6.1)^18^, against the eukaryota_odb12, viridiplantae_odb12, embrophyta_odb12, and liliopsida_odb12 databases. The *K*-mer (21-mers) completeness and assembly consensus quality value (QV) were estimated using Merqury (v1.4.1)^17^. The QV is a log-scaled probability of errors for consensus base calls; the higher the QV values, the more accurate the consensus base calls, with a QV value at 30 indicating 99.9% accuracy and 40 representing 99.99% accuracy^17^. The regional and structural assembly quality indicator (R-AQI and S-AQI) were assessed using the Clipping information for Revealing Assembly Quality (CRAQ, v1.0.9) algorithm ^16^; the higher the AQI scores, the more accurate the genome assembly^16^. The long terminal repeat retrotransposons (LTR-RTs) were first annotated using LTRharvest (v1.6.1)^40^ with the parameters ‘-maxlenltr 6000 -mindistltr 1500 -maxdistltr 25000 -mintsd 5-maxtsd 5 -vic 10 -similar 85’ and LTR_FINDER_parallel (v1.2)^41^ with the parameters ‘-size 1000000-time 300’. Then the LTR Assembly Index (LAI)^42^ was calculated using LTR_retriever (v2.9.5)^43^ with the parameter ‘-u 0.5e-8’. The telomere and centromeric repeat identification were conducted using quartet (v1.2.5) using default parameter settings^44^.

### Noncoding RNA and repetitive sequence identification

Noncoding RNAs were predicted using tRNAscan-SE (v2.0.9)^45^ and Infernal software (v1.1.4)^46^ by searching the database Rfam (v14.10). The extensive *de-novo* TE annotator (EDTA) (v2.1.0)^47^—that encompasses LTR_FINDER, LTRharvest, LTR_retriver, Generic Repeat Finder, TIR-Learner, HelitronScanner, RepeatModeler and RepeatMasker— was used to identify, annotate and classify repetitive elements of the genome, with parameters ‘--anno 1 --sensitive 1 --evaluate 1 --u 0.5e-8 --curatedlib ${SINE_LINE_lib}’. The library of short interspersed elements (SINEs) and long interspersed nuclear elements (LINEs) was downloaded from SINEBase (https://sines.eimb.ru/) as a compensation for the reference sequences. The repetitive elements identified by EDTA included long terminal repeat (LTR) retrotransposons (copia, gypsy and unknown ones) and non-LTR retrotransposons (LINE_element and unknown ones), terminal inverted repeat (TIR) transposons (CACTA, Mutator, PIF_Harbinger, Tc1_mariner and hAT), *Helitron* and other interspersed repeats. Subsequently, the DeepTE (version updated on Oct 22^th^, 2022) script^48^ was used to reclassify the TEs that were initially annotated as ‘LTR/unknown’ by EDTA into copia, gypsy and unknown classes, then the output file was fed back into EDTA to enhance the comprehensiveness of TE annotation. To calculate LTR insertion time, we set the mutation rate per generation (μ) to 0.5×10^-8^, based on the assumption that the generation times of orchids are four years^49^. The resulting hard-masked genome was converted into a soft-masked genome, which was used in the subsequent analysis. The tandem repeat sequences in the *P.* sa genome were identified using Tandem Repeats Finder (TRF, v4.09)^50^ with the parameters ‘2 7 7 80 10 50 2000 -d -h’. Then the proportion of tandem repeats in the genome was calculated using the script buildRepeatSummary.pl (v1.36) with default parameters^51^.

### Transcriptome sequencing

Total RNA was extracted for roots, leaves, stems and flower buds with mixed stages (one duplicate) and other tissues at different development stages (three replicates, **Table S6**). The library preparation and PCR amplification were performed according to the manufacturer’s instructions, and the RNA libraries were sequenced on the Illumina Novaseq 6000 platform and 150 bp paired-end reads were generated. Clean data were further obtained by removing reads containing adapters, reads containing poly-N and low quality reads from raw data, using fastp (v0.24.1)^52^ with parameters: ‘-g -q 5 -u 50 -n 15 -l 150 --overlap_diff_limit 1 --overlap_diff_percent_limit 10’.

### Protein-coding gene prediction

Protein-coding genes were predicted by integrating predictions based on sequence homology and transcriptome, and the *ab initio* predictions using the soft-masked genome.

#### 1. Homology-based prediction

Genome assembly and General Feature Format (GFF) files of six plant species were used for homology-based prediction with Gene Model Mapper (GeMoMa, v1.9)^53^. The following species were included: *Arabidopsis thaliana* (TAIR10.1, downloaded from NCBI), *Oryza sativa* Japonica Group (IRGSP_1.0, from Ensembl Plant), *Asparagus officinalis* (Aspof.V1, from Ensembl Plant), *Apostasia shenzhenica* (GCA_002786265.1, from NCBI)^54^, *Dendrobium catenatum* (GCF_001605985.2, from NCBI)^55^, and *Phalaenopsis equestris* (GCF_001263595.1, from NCBI)^14^. The GeMoMa parameters used were ‘AnnotationFinalizer.r=NO p=false o=true tblastn=false’.

#### 2. Transcriptome-based prediction

Clean RNA-seq reads from roots, leaves, stems, and mixed-stage flower buds were aligned to *P. sa* genome using HISAT2 (hierarchical indexing for spliced alignment of transcripts, v2.2.1)^56^. The alignments were processed by StringTie (v2.2.1)^57^ to reconstruct genome-guided transcripts. Potential open reading frames (ORFs, ≥100 amino acids) were identified from these transcripts using TransDecoder (v5.7.1) (https://github.com/TransDecoder). A genome-guided transcriptome was assembled using Trinity (v2.15.1)^58^. Both the StringTie output and the Trinity assembly were then analyzed by the Program to Assemble Spliced Alignments (PASA, v2.5.3)^59^ with the parameters ‘--ALIGNERS blat,minimap2 --TRANSDECODER’ to generate transcriptome-supported gene models.

#### 3. *Ab initio* prediction

The BRAKER3 pipeline (v3.0.3, incorporating Augustus and Genemark)^60^, GeneID (2021 version)^61^ and GlimmerHMM (2014 version)^62^ were used for the *ab initio* gene prediction, leveraging the transcriptome evidence as a guide.

#### 4. Gene model integration and filtering

The EvidenceModeler (v2.1.0)^63,64^ was used to integrate predictions from the above methods into consensus gene structures. The following weights were assigned: GeneID (1), GlimmerHMM (1), GeneMark (2), Augustus (4), GeMoMa (7), StringTie (5), and PASA_transdecoder (10).

To refine the gene models, predicted protein sequences were searched against the Transposon Protein Database (https://github.com/NBISweden/TransposonPSI/blob/master/transposon_ORF_lib/transposon_db.pep) using TransposonPSI (v1.0.0), which employs PSI-BLAST to identify TE-related sequences^65^ with default parameters. Proteins with > 30% similarity and > 60% coverage to known TE sequences were removed. In addition, gene models with incomplete ORFs or ORFs ≤ 100 amino acids were excluded, if they met all the three following conditions: lacking RNA-seq expression support, having < 50% coverage overlap with GeMoMa models, and lacking functional annotation by InterProScan (v5.67-99.0)^66^.

#### 5. Model optimization with extended RNA-seq data

The resulting gene models (v1.0) from above steps were further supplemented using additional RNA-seq data from other 28 samples (**Table S6**). First, to avoid potential mapping errors between tandem duplicates with high sequence similarities, RNA-seq reads were first mapped to the v1.0 transcriptome, and unmapped reads were further mapped to the genome, using TopHat2 (v2.1.1)^67^ with the parameter ‘--max-multihits 1’. Next, individual bam files were used to assemble transcripts (including alternative splicing isoforms) using StringTie. The resulting GTF files from all samples were merged using StringTie to create a comprehensive transcript annotation. TransDecoder was used to infer the ORFs, producing a GFF file (v2.0) that included alternative splicing and five prime untranslated regions (5’UTR) and 3’UTR information. Finally, the v2.0 annotations were compared with and used to correct the v1.0 annotations, prioritizing v2.0 evidence in overlapping regions. This process resulted in a final set of 93,708 protein-coding genes.

### Protein-coding gene functional annotation

*P.* sa protein sequences were first searched against the eggNOG database (v5.0.2)^68^ using eggNOG-mapper (v2.1.12)^69^, with parameters ‘-m diamond --seed_ortholog_evalue 1e-5 --tax_scope Viridiplantae’, for annotations including Gene Ontology (GO), Kyoto Encyclopedia of Genes and Genomes (KEGG) Orthology (KO), and Clusters of Orthologous Groups (COG). In order to obtain the functional description of related protein-coding genes in model plant species, protein sequences of *P.* sa genome were searched against the UniProt database (uniprot_sprot_plants.fasta that contains manually annotated protein sequences from sequenced plant genomes, updated on March 27^th^, 2024, https://www.uniprot.org/) and protein sequences from Arabidopsis (Araport11_pep_20240409, downloaded from The Arabidopsis Information Resource) using BLASTP (v2.12.0+)^70^ with parameters ‘-max_hsps 1 -max_target_seqs 1 -evalue 1e-5’. Furthermore, *P*. sa protein sequences were searched against databases such as Pfam, SMART, CDD, Superfamily and others, using InterProScan (v5.67-99.0) with parameters ‘-goterms -iprlookup’. Gene Functional Annotation for Plants (GFAP, v3.1)^71^ was used to annotate GO, KEGG and Pfam items by searching *P*. sa protein sequences against a plant-specific database with the parameter ‘-awd psd -go -kegg -pfam’, and to obtain homology-based annotations by searching against the plant-specific NCBI non-redundant (nr) database with the parameter ‘-awd nr’. Lastly, protein sequences were searched against the Reference Protein Sequence Dataset (RPSD, release_2024_07_31) using Ensemble Enzyme Prediction Pipeline (E2P2, v5)^72^ with default parameters for the enzyme function annotation. Transcription factors in *P.* sa genome were annotated in the Plant Transcription Factor Database (PlantTFDB)^73^.

### Homoeologous gene identification

The Synteny and Rearrangement Identifier (SyRI, v1.7.0)^74^ and MCScanX v1.0.0^75^ were used to conduct synteny analysis among different HChrs with default parameter settings, based on genomic and protein sequences, respectively. Later on, the homoeologous genes were identified with in-house scripts (see **Code availability**) using the outputs of SyRI preferentially and MCScanX supplementarily, to avoid improper assignment of multiple MCScanX short syntenic blocks between HChrs.

### Transcriptome analysis

RNA-seq reads were mapped to the transcriptome and genome using Tophat2 (v2.1.1)^67^, with parameter ‘--max-multihits 1’, using indexes from both the genome and the final transcriptome. Read counts for protein-coding genes were obtained using htseq-count (v2.0.5)^76^ and TPM was calculated using the “calculateTPM” function in R package scater (v1.0.4)^77^. The TPM values were normalized across samples using quantile normalization, followed by averaging across the three biological replicates to obtain a mean TPM value for each tissue.

To categorize genes according to their expression levels, we defined four thresholds based on TPM values: 1) TPM = 0 indicating no expression; 2) 0 < TPM ≤ 2 indicating low expression; 3) 2 < TPM ≤ 10 indicating moderate expression; 4) TPM > 10 indicating high expression. The threshold of TPM = 10 distinguishing high expression was determined by modeling the observed gene expression data as a mixed distribution. This model comprises a normal distribution, representing true biological expression, and an exponentially decaying distribution, representing technical noise or background. These two components were fitted using the nls function from the R “stats” package^78^. The value of TPM = 10 corresponds approximately to the 50^th^ percentile of the fitted normal distribution, suggesting that roughly half of the genuine gene expression occurs below this level.

### Weighted gene co-expression network analysis (WGCNA)

Co-expression analysis for HG type 1 genes that were highly expressed (TPM > 10) in ≥ 1 tissue and had no non-self blast hist in *P.* sa genome was performed using WGCNA (v1.72.1)^79^, under a signed network model, with a minimum module size of 10.

### Gene tree building

To build a robust phylogeny tree for *MYB2*, *MYB11* and *MYB12* genes, coding sequences (CDS) were obtained from multiple orchid species—*P. aphrodite* (*P. ap*), *P. equestris* (*P. eq*), *Apostasia shenzhenica* (*A. sh*), *Cremastra appendiculata* (*C. ap*), *Cymbidium sinense* (*C. si*), *Dendrobium chrysotoxum* (*D. ch*), *Gastrodia elata* (*G. el*), *Platanthera guangdongensis* (*P. gu*), *Vanilla planifolia* (*V. pl*)—via the Orchid Multiomics Database (http://orchidcomics.com/, access on June 11^th^, 2025)^80^. For each query gene, the top 20 BLAST hits per species (top 80 hits for *P.* sa) were extracted, translated into amino acid sequences, and aligned using MEGA v7.0^81^. The resulting amino acid alignment was manually curated to retain only reliably alignable regions, which were then converted back to a corresponding CDS alignment. The best-fit nucleotide substitution model (GTR+I+G4) was selected using ModelTest-NG (v0.1.7)^82^. Finally, a maximum likelihood phylogenetic tree was constructed with RAxML-NG (v1.2.2)^83^, employing 1000 bootstrap replicates.

The subclass of floral organ identity genes was determined by phylogenetic analysis. CDS of MIKC-type MADS-box genes from *P.* sa and other orchids (**Fig. S10**) were translated into amino acid sequences. These sequences were aligned using MEGA with manual adjustment. A phylogeny was inferred using the ‘JTT+I+G4’ model with 1000 bootstrap iterations.

### Genetic variant calling

Total DNA was extracted for flower stalks of 46 H80 (*P*. ‘1071-13’ × *P*. ‘CH1804-01’) individuals. Sequencing services of partial H80 individuals were provided by BGI Genomics (Shenzhen, China), and the others were provided by Novogene Co., Ltd (Beijing, China). All whole-genome resequencing in this study was performed on the DNBSEQ-T7 platform using a 150-bp paired-end strategy, generating ∼60 Gb of data per sample. Raw reads were processed using fastp (v0.24.1)^52^ with parameters ‘-g -q 5 -u 50 -n 15 -l 150 --overlap_diff_limit 1 --overlap_diff_percent_limit 10’. Clean reads were aligned to the *P*. sa genome using BWA-MEM algorithm in Burrow-Wheeler Aligner (BWA, 0.7.18-r1243-dirty)^84^ with default settings, followed by sorting with SAMtools (v1.21)^85^. Duplicate reads were marked using the MarkDuplicates module in the Genome Analysis Toolkit (GATK, v4.6.1.0)^86^.

Population-level SNP and indel calling was conducted using GATK. First, variants were called per sample with HaplotypeCaller using ‘-ERC GVCF -ploidy 4’. Individual gVCF files were merged with CombineGVCFs and jointly genotyped with GenotypeGVCFs. SNPs and indels were separated using SelectVariants and filtered with VariantFiltration using the following criteria: for SNPs, ‘QD < 2.0 || MQ < 40.0 || FS > 60.0 || SOR > 3.0 || MQRankSum < -12.5 || ReadPosRankSum < -8.0’; for indels, ‘QD < 2.0 || FS > 200.0 || MQ < 40.0 || SOR > 10.0 || ReadPosRankSum < -20.0’. Multiallelic sites were split into biallelic records using BCFtools norm (v1.21)^85^ with ‘--multiallelics-both’.

SVs were called and genotyped using smoove (v0.2.8)^87^ with default parameters. CNVs were detected with CNVpytor (v1.3.2)^88^ using a bin size of 100. Finally, all variants (SNPs, indels, SVs and CNVs) were annotated using ANNOVAR (v2020-06-07)^89^.

### Genome-wide association study (GWAS) analysis

GWAS was performed using the R package GWASpoly (v2.13)^90^ based on the Q + K mixed model. First, the VCF files were transformed into dosage files using VCF2dosage function with parameters ‘-ploidy 4 -min.DP 8 -max.missing 0.5 -min.minor 3’. To control for population structure, the covariance matrix for polygenic effect was calculated using the set.K function with the parameter ‘LOCO=TRUE’, where a different covariance matrix was calculated for each chromosome according to markers from all other chromosomes^91^. During this step, missing data were imputed using the k-Nearest Neighbors approach implemented by the KNNImputer function from the sklearn.impute module^92^. The imputation was performed five times with different k values (2, 5, 10, 15, and 20), and the average of the five imputed values was used. The set.params function was used to set the threshold of minor allele frequency with the parameter ‘geno.freq = 1 - 5/N, MAF=0.05’. The GWASpoly function was then used to assess the significance of each marker associated with target trait variation under the additive model and two simplex dominance models (1-dom-ref and 1-dom-alt) hypothesis. In the additive model, the marker effect scales linearly with the alternate allele dosage. For the simplex dominance models, a single copy of the dominant allele (reference or alternate allele) is sufficient for the complete dominance phenotype, thereby all heterozygotes possessing this allele exhibit equivalent marker effects. Two significance thresholds were adopted: *p*-value<0.05/variant counts and *p*-value<0.01/variant counts, according to the Bonferroni adjustment approach. Lastly, the CMplot (v4.5.1)^93^ package was used to generate the Manhattan plots. For better visualization, -log_10_(*p*) values greater than 30 were truncated to 30 in the Manhattan plots.

### Virus-induced gene silencing

Seedlings of *P.* ‘Jinbianlinglong’ (*P.* ji) were cultivated in a growth room under a 14-h light/10-h dark photoperiod, with a light intensity of 220 µmol·m^-2^·s^-1^ and a temperature of 25–27 ℃. Partial sequences of *PjPDS* and *PjMYBx1* were introduced into the TRV2-based pYL156 vector, as illustrated in **Fig. S21**. The resulting TRV2 plasmids were transformed into *Agrobacterium tumefaciens* strain GV3101. Bacterial cultures containing either TRV2-*PDS* or TRV2-*PDS*-*MYBx1*, and cultures containing the TRV1 plasmids were incubated separately. When the OD_600_ reached approximately 2.0, the cells were harvested and resuspended in an induction buffer (10 mM morpholineethanesulfonic acid, 10 mM MgCl_2_, and 200 uM acetosyringone) to a final OD_600_ of 2.0. Each TRV2 suspension was mixed with an equal volume of the TRV1 suspension, and Silwet L-77 was added to a final concentration of 0.005%. The mixture was incubated statically at room temperature for 3 h.

The resulting *Agrobacterium* mixture was infiltrated into the newly developed inflorescence stalk and its nearest leaf. After infiltration, plants were kept in darkness for 48 h before being returned to the growth room. Plants were monitored until phenotypic changes in the flowers became evident.

To confirm the silencing of target genes in plants showing phenotypic changes, total RNA was extracted from the corresponding floral tissues using the FastPure Universal Plant Total RNA Isolation Kit (Vazyme, Nanjing, China). First-strand cDNA was synthesized using MightyScript Plus First Strand cDNA Synthesis Master Mix (gDNA digester) (Sangon Biotech, Shanghai, China). Quantitative reverse-transcription PCR (qRT-PCR) was performed using SupRealQ Ultra Hunter SYBR qPCR Master Mix (U+) (Vazyme). The 2^−ΔΔCt^ method was employed to calculate the relative expression of target and related genes, with *PeActin* serving as the internal control. Primer sequences for VIGS and qRT-PCR were listed in **Table S11.**

## Data availability

The raw sequencing data used in this study have been submitted to the NCBI database under BioProject PRJNA1143085 (will be released upon publication), including PacBio HiFi, Hi-C and transcriptome sequences for *P.* sa, and whole genome sequences for 46 H80 individuals. The *P.* sa genome assembly, annotation, homoeologous gene table, and expression matrix for all protein-coding genes will be released in our database (http://www.phaldb.cn/) upon publication.

## Code availability

Custom scripts and codes used in this study are provided at GitHub (https://github.com/peipeiwang6/Manuscript/tree/main/2025_Psa_genome).

## Acknowledgement

We thank Dr. Huiqi Zhao for sharing the protocol for karyotype analysis, Prof. Zhiqiang Wu, Prof. Xingtan Zhang and Dr. Caiyao Zhao for their helpful suggestions on genome sequencing and assembling, Dr. Xukun Li for advice on anatomical photography and other experiments, Fu Chen for providing seedlings of *P.* ‘Jinbianlinglong’. The English language of this manuscript was partially polished using DeepSeek, while all original content was written by the authors and the revised text was adopted following author review.

This work was supported by the National Natural Science Foundation of China (32370241), the Scientific Research Foundation of Kunpeng Institute of Modern Agriculture at Foshan (KIMAQD2022003), and the collaborative project (HXQS2024009 from Prof. Jue Ruan) to P.W.; National Key R&D Program of China (2023YFD1600505) to R.J.; Project supported by the Foundation for Innovative Research Groups of the National Natural Science Foundation of China (32221001) to H.K.; Chinese Universities Scientific Fund (2452024391) to R.Z.

## Author contributions

PW, RJ, HK and RZ conceived and designed the study. PW, XZ, LZ, ZL, ZW, FW, YJ, YZ, and LW collected the plant materials of *P.* sa for genome and transcriptome sequencing; RJ conducted the hybridization and provided the plant materials; RJ, LW, YZ and XC conducted the karyotype analysis; PW, LZ, HH and LLan (Lan Lan) assembled the genome; LZ and PW conducted the genome annotation; LZ conducted the functional annotation for protein-coding genes; LZ, FM and PW conducted the genome assembly quality assessment and syntenic analysis; PW and FW conducted the transcriptome analysis; PW, XZ and FW conducted the gene tree building; XZ conducted the genetic variant calling and GWAS analyses; ZW, LLin (Letian Lin) and ZL conducted the VIGS experiments; LLin performed the qRT-PCR; PW conducted all the other analyses; PW, RJ, XZ, LZ, LW, ZW and LLin wrote the draft with contributions from all the other authors. All authors read and approved the final manuscript.

## Supplementary figure legends

**Fig. S1 Phenotypes of H80 individuals.** Each panel shows an individual plant, with its classifications for the P/A of stripe, background pink color and spot/patch, and lip morphology provided on the top-left, top-right, bottom-left and bottom-right corners per panel, respectively. Red arrowheads indicate stripes, while blue ones indicate venation-associated shadow.

**Fig. S2 Genome survey of *P.* sa and karyotype analysis for H80.** (**A,B**) *k*-mer coverage distribution of *P.* sa sequencing reads, under the assumptions that *P.* sa is a triploid (**A**) or tetraploid (**B**). (**C**) *k*-mer coverage analysis indicates *P.* sa is an allo-triploid. (**D**) Karyotype analysis reveals 70 chromosomes per cell in *P.* sa. (**E**) Inferred long terminal repeats (LTRs) insertion time. (**F**) Karyotype analysis reveals 64 chromosomes per cell in H80 individuals.

**Fig. S3 Subgenome assignments of 70 *P*. sa chromosomes according to subgenome-specific *k*-mers.** From outer to inner circles (1-9): (1) subgenome assignments; (2) significant enrichment of subgenome-specific *k*-mers; (3) normalized proportion of subgenome-specific *k*-mers; (4-7) count of subgenome-specific *k*-mers for four sets, respectively; (8) density of long terminal repeat retrotransposons (LTR-RTs), with gray indicating nonspecific LTR-RTs; (9) homoeologous blocks.

**Fig. S4 Expression profiles of *FAR-RED IMPAIRED RESPONSE 1 (FAR1*) genes in *P.* sa.** *FAR1* genes are grouped into different HG types, as illustrated in **Fig. 2A**. Color scale in the heatmap indicates log-transformed Transcripts Per Kilobase (TPM) values, where blue and light blue indicate original TPM=0 and TPM<2, respectively.

**Fig. S5 Expression profiles of potential *de novo* new genes.** (**A-G**) Expression profiles of seven co-expression modules inferred from WGCNA analysis (**Methods**) for type 1 genes that were highly expressed (TPM>10) in ≥ 1 tissue and had no non-self blastp hists in *P.* sa genome. *Z*-scores of TPM values among all the tissues are shown. Example genes that were preferentially expressed in different tissues are indicated.

**Fig. S6 Protein sequence alignment (A) and expression (B) of *PsAGL6-2* HGs.** HGs with truncated sequences were indicated with green shadow.

**Fig. S7 Manhattan plot for GWAS of lip morphology in the H80 population using HChr set a as the reference.** (**A-D**) GWAS results when SNPs, indels, SVs and CNVs were used, respectively, with the additive marker-effect model. Horizontal solid line and dashed line indicate -log_10_ (0.01/variant counts) and -log_10_ (0.05/variant counts) (**Methods**), respectively; red and orange dots denote variants significantly associated with the lip morphology variation; blue dots represent variants located within or adjacent (±2 kb) to the tandem duplicate cluster of *PsAGL6-2a* genes.

**Fig. S8 Manhattan plot for GWAS of lip morphology in H80 using HChr set b as the reference.** (**A-D**) GWAS results when SNPs, indels, SVs and CNVs were used, respectively, with the additive marker-effect model. Red and orange dots denote variants significantly associated with the lip morphology variation; blue dots represent variants located within or adjacent (±2 kb) to the tandem duplicate cluster of *PsAGL6-2b* genes.

**Fig. S9 Manhattan plot for GWAS of lip morphology in H80 using HChr set c as the reference.** (**A-D**) GWAS results when SNPs, indels, SVs and CNVs were used, respectively, with the additive marker-effect model. Red and orange dots denote variants significantly associated with the lip morphology variation; blue dots represent variants located within or adjacent (±2 kb) to the tandem duplicate cluster of *PsAGL6-2c* genes.

**Fig. S10 Gene tree for MIKC-type MADS-box genes in orchids.** NCBI accession ID (such as JL017169) was listed between gene name (such as *CeMADS32*) and the reference (such as 2021 Ai).

**Fig. S11 Manhattan plot for GWAS of stripe P/A in H80 using the additive marker-effect model.** (**A-D**) GWAS results when SNPs, indels, SVs and CNVs that were called by referring to HChr set a were used, respectively. (**E-H**) GWAS results when SNPs, indels, SVs and CNVs that were called by referring to HChr set b were used. Red and orange dots denote variants significantly associated with the P/A variation of stripe; blue dots represent variants located within or adjacent (±2 kb) to the tandem duplicate cluster of *PsMYB12a* genes.

**Fig. S12 IGV images showing reads of seven H80 individuals with or without stripes that were mapped onto the *PsMYB12* gene-containing regions.** Red arrowheads indicate stripes, while blue ones indicate venation-associated shadow in individuals which had no intact reads mapped onto *PsMYB12* genes.

**Fig. S13 Evolution of color patterning-related *MYB* genes.** (**A**) Gene tree of *MYB* genes from orchids with genome assemblies. (**B**) Evolutionary history of stripes on perianths in the genus *Phalaneopsis*. The phylogeny of *Phalaenopsis* species were modified from our previous review paper ^28^. The red star indicates the origination of stripes in the most recent common ancestor of *P. equestris* and *P. lindenii*. Three flower images show the venation-associated distribution of spots in other *Phalaenopsis* species. (**C**) Integrative Genomics Viewer images showing reads of *Vanda* ‘Pakchong Blue’ that were mapped onto genic regions of two *PsMYB12* genes as examples.

**Fig. S14 Homoeologous gene variation in sequence and expression of *PsMYB2* and *PsMYB11*.** (**A**) Protein sequence alignment of *PsMYB2* HGs. (**B**) HG-specific expression of *PsMYB2* in *P*. sa floral organs (upper) and the lips of selected H80 individuals (lower), with values in the table indicating TPM values. (**C,D**) Protein sequence and promoter DNA sequence (1 Kb upstream) alignments of *PsMYB11* HGs, respectively. (**E**) HG-specific expression of *PsMYB11* in *P*. sa floral organs (upper) and the lips of selected H80 individuals (lower). The HG showing high sequence divergence with others was indicated using green shadow.

**Fig. S15 Manhattan plot for GWAS of background color in H80 using the additive marker-effect model.** (**A-D**) GWAS results when SNPs, indels, SVs and CNVs that were called by referring to HChr set a were used, respectively. Red and orange dots denote variants significantly associated with the P/A variation of pink color; blue dots represent variants located within or adjacent (±2 kb) to *PsMYB2a*.

**Fig. S16 Manhattan plot for GWAS of background color in H80 using the 1-dom-ref marker-effect model.** (**A-D**) GWAS results when SNPs, indels, SVs and CNVs that were called by referring to HChr set b were used, respectively. Red dots denote variants significantly associated with the P/A variation of pink color; blue dots represent variants located within or adjacent (±2 kb) to *PsMYB2b*.

**Fig. S17 Manhattan plot for GWAS of background color in H80 using the 1-dom-alt marker-effect model.** (**A-D**) GWAS results when SNPs, indels, SVs and CNVs that were called by referring to HChr set b were used, respectively. Red and orange dots denote variants significantly associated with the P/A variation of pink color; blue dots represent variants located within or adjacent (±2 kb) to *PsMYB2b*.

**Fig. S18 Manhattan plot for GWAS of patch P/A in H80 using HChr set a as the reference.** (**A-D**) GWAS results when SNPs, indels, SVs and CNVs were used, respectively, with the additive marker-effect model. Red and orange dots denote variants significantly associated with the P/A variation of patch; blue dots represent variants located within or adjacent (±2 kb) to *PsMYB11a*.

**Fig. S19 Manhattan plot for GWAS of patch P/A in H80 using HChr set b as the reference.** (**A-D**) GWAS results when SNPs, indels, SVs and CNVs were used, respectively, with the additive marker-effect model. Red and orange dots denote variants significantly associated with the P/A variation of patch; blue dots represent variants located within or adjacent (±2 kb) to *PsMYB11b*.

**Fig. S20 Manhattan plot for GWAS of patch P/A in H80 using HChr set c as the reference.** (**A-D**) GWAS results when SNPs, indels, SVs and CNVs were used, respectively, with the additive marker-effect model. Red and orange dots denote variants significantly associated with the P/A variation of patch; blue dots represent variants located within or adjacent (±2 kb) to the tandem duplicate cluster of *PsMYB11c* genes.

**Fig. S21 *PjPDS* and *PjMYBx1* sequences and the VIGS constructions.** (**A**) CDS and 3’UTR sequence alignment for four *PjPDS* HGs. (**B**) CDS sequence alignment for four *PjMYBx1* HGs. (**C**) Diagrams showing the structure of TRV2 plasmid PYL156, and the insertion locations of *PjPDS* and *PjMYBx1* sequences. *PjPDS_1* and *PjPDS_2* are sequences from two *PjPDS* HGs marked with green shadow.

## Supplementary tables

**Table S1** Comparison of basic statistics for genome assembly and annotation of three *Phalaenopsis* species

**Table S2** Statistics of repetitive sequences in *P.* sa genome

**Table S3** Statistics of *P.* sa genes and the functional annotations

**Table S4** Statistics of noncoding RNAs in *P*. sa genome

**Table S5** Structural variation summary

**Table S6** Information for RNA-seq samples

**Table S7** IDs for genes mentioned in the manuscript

**Table S8** Allele read ratios of two SNPs in H80 individuals with different lip morphology

**Table S9** Association between the presence/absence of stripes and *PsMYB12* genes in 46 H80 individuals

**Table S10** Association between *PsMYB2b*-type sequences with background colors

**Table S11** Primer sequences used for VIGS and qRT-PCR

