## Supplementary material for "Haplotype-resolved genome assembly advances genetic understanding of trait diversity in *Phalaenopsis* orchids": Fig. S1-S21

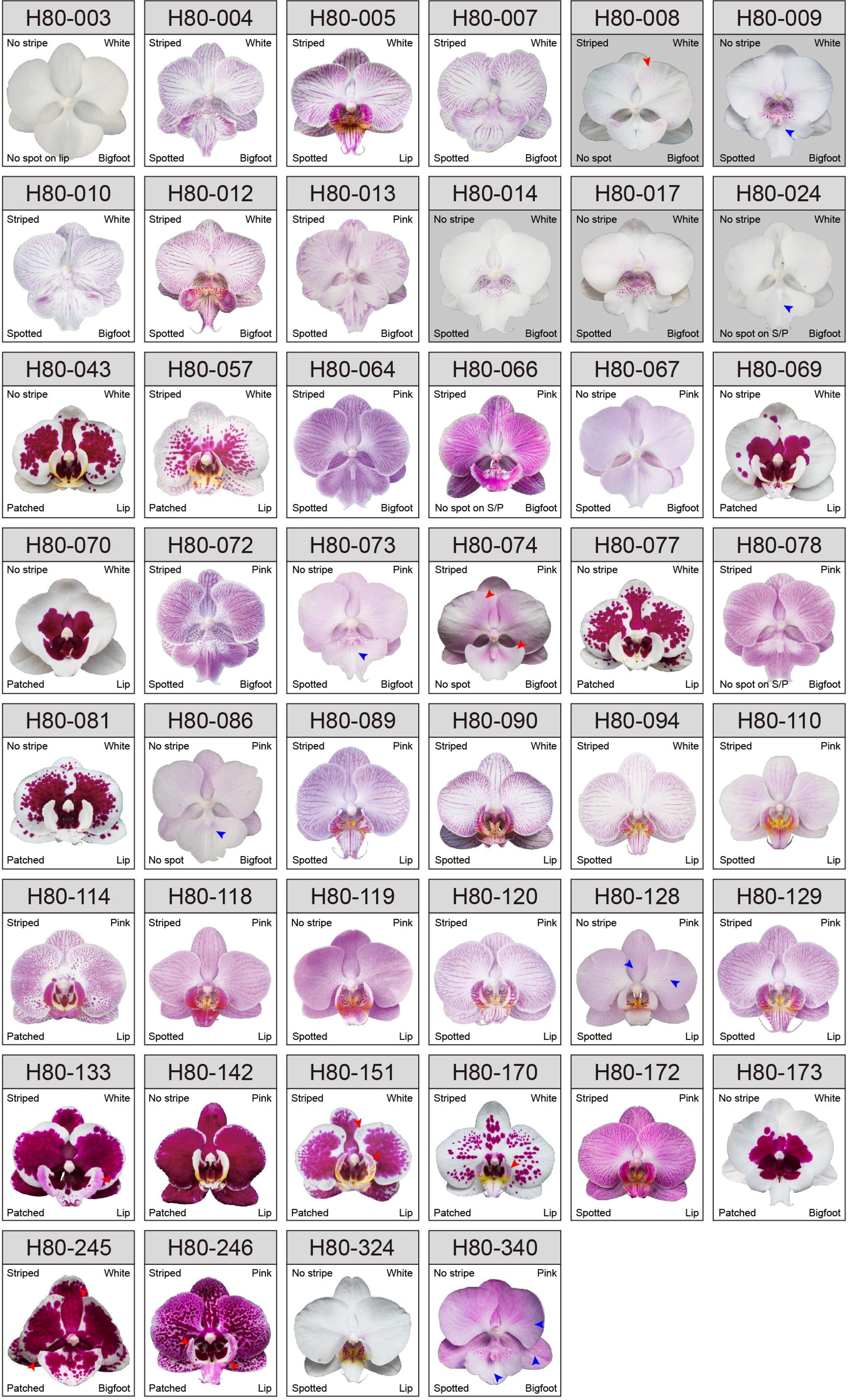


**Fig. S1 Phenotypes of H80 individuals.** Each panel shows an individual plant, with its classifications for the P/A of stripe, background pink color and spot/patch, and lip morphology provided on the top-left, top-right, bottom-left and bottom-right corners per panel, respectively. Red arrowheads indicate stripes, while blue ones indicate venation-associated shadow.

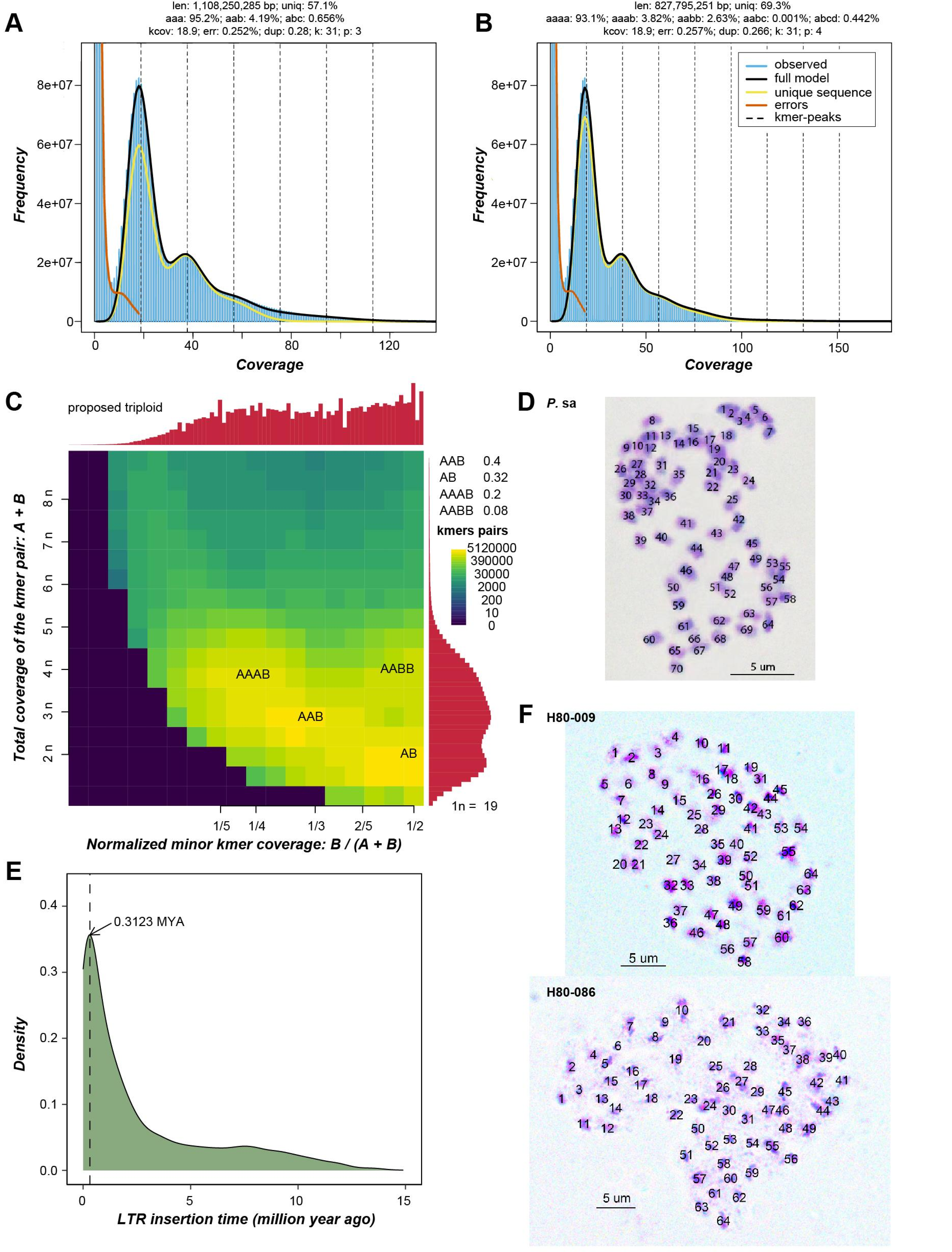


**Fig. S2 Genome survey of *P.* sa and karyotype analysis for H80.** (**A,B**) *k*-mer coverage distribution of *P*. sa sequencing reads, under the assumptions that *P*. sa is a triploid (**A**) or tetraploid (**B**). (**C**) *k*-mer coverage analysis indicates *P*. sa is an allo-triploid. (**D**) Karyotype analysis reveals 70 chromosomes per cell in *P*. sa. (**E**) Inferred long terminal repeats (LTRs) insertion time. (**F**) Karyotype analysis reveals 64 chromosomes per cell in H80 individuals.

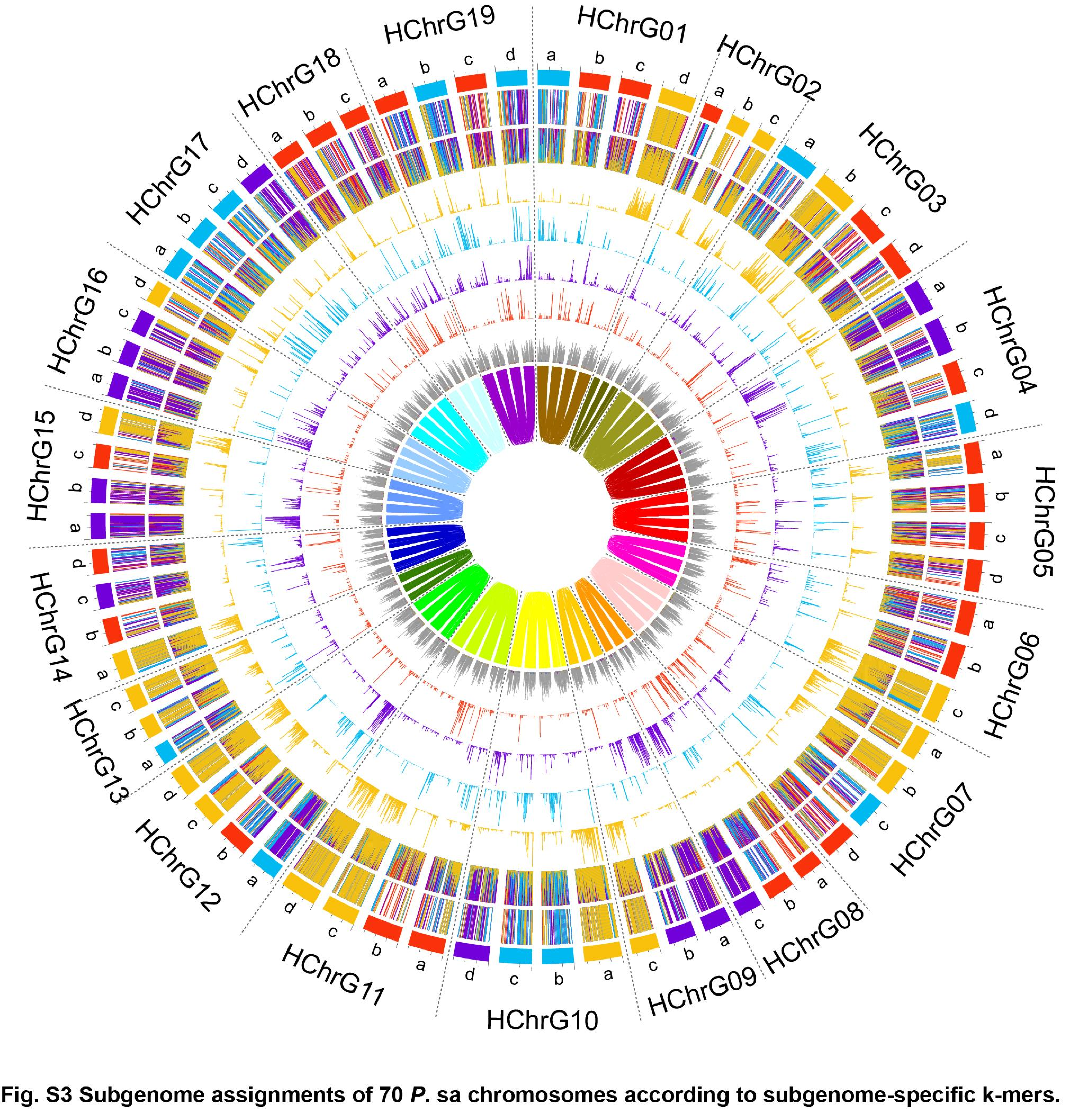


**Fig. S3 Subgenome assignments of 70 *P*. sa chromosomes according to subgenome-specific k-mers.** From outer to inner circles (1-9): (1) subgenome assignments; (2) significant enrichment of subgenome-specific *k*-mers; (3) normalized proportion of subgenome-specific *k*-mers; (4-7) count of subgenome-specific *k*-mers for four sets, respectively; (8) density of long terminal repeat retrotransposons (LTR-RTs), with gray indicating nonspecific LTR-RTs; (9) homoeologous blocks.

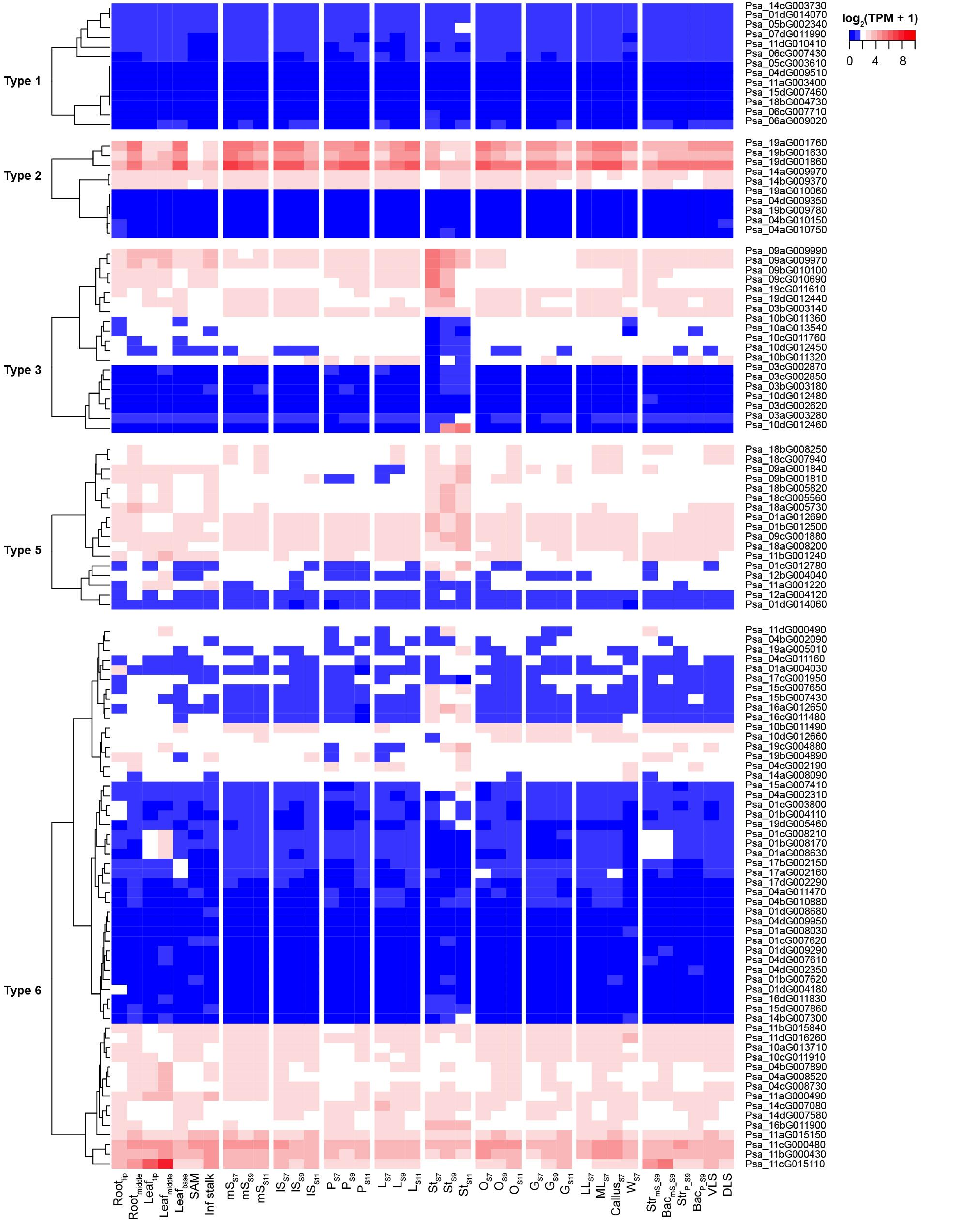


**Fig. S4 Expression profiles of *FAR-RED IMPAIRED RESPONSE 1* (*FAR1*) genes in *P*. sa.** *FAR1* genes are grouped into different HG types, as illustrated in **Fig. 2A**. Color scale in the heatmap indicates log-transformed Transcripts Per Kilobase (TPM) values, where blue and light blue indicate original TPM=0 and TPM<2, respectively.

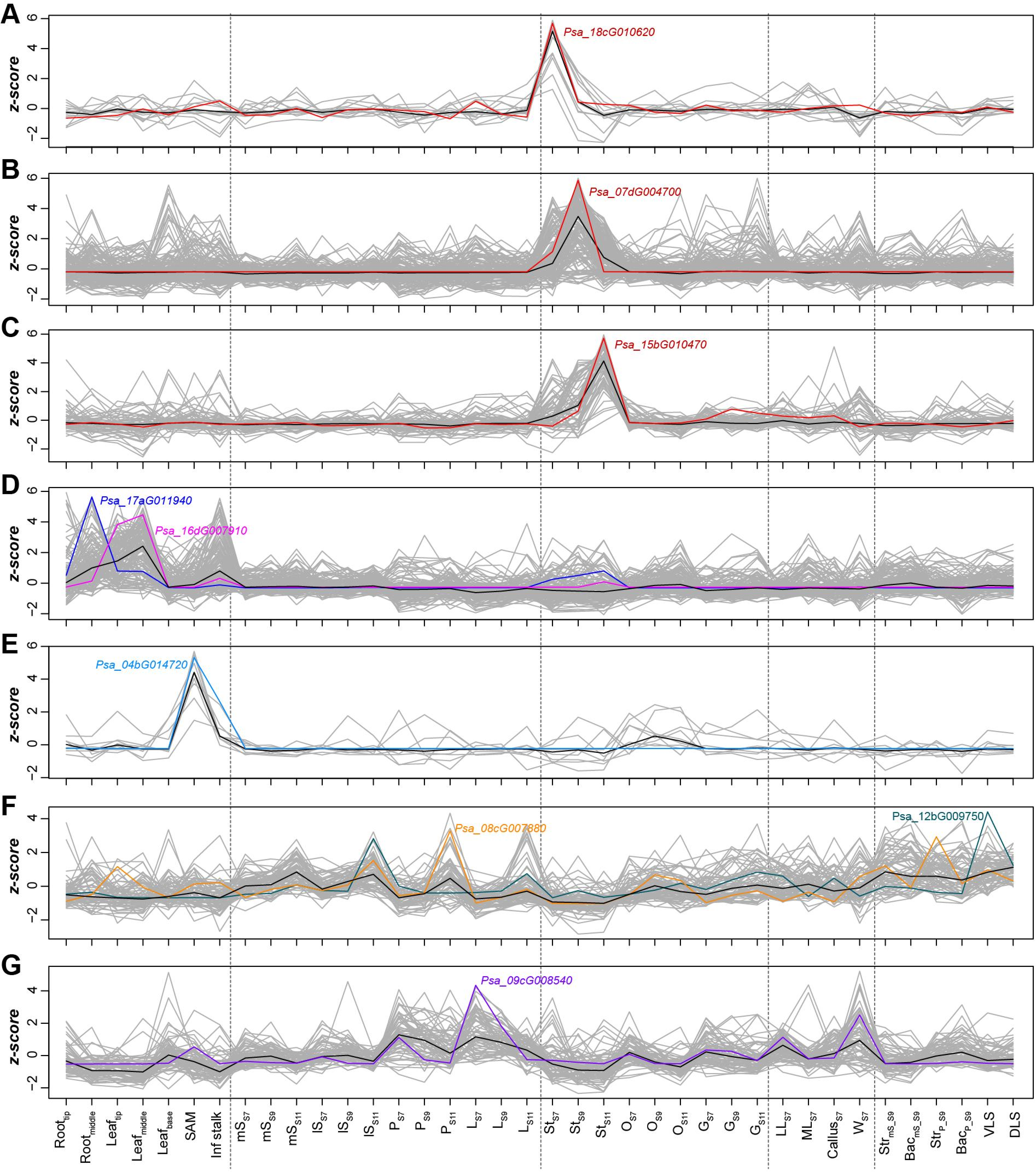


**Fig. S5 Expression profiles of potential *de novo* new genes.** (**A-G**) Expression profiles of seven co-expression modules inferred from WGCNA analysis (**Methods**) for type 1 genes that were highly expressed (TPM>10) in ≥ 1 tissue and had no non-self blastp hists in P. sa genome. *Z*-scores of TPM values among all the tissues are shown. Example genes that were preferentially expressed in different tissues are indicated.

**A**

```
Consensus     MGRGRVELKRIENKI NRQVTFSKRRNGIMK KAYELSVLCDAEIAL IIFSSRGKLFEFGSP DITKTLERYQRCTFT PQTIHPNDHETLNWY QELSKLKAKYESLQR SQRHLLGEDLDLLNL KELQQLERQLESSLS QARQKRTQIMLDQME 150
Psa_09aG004230 --------------- --------------- --------------- --------------- --------------- --------------- --------------- --------------- --------------- ----MQ.........  11
Psa_09aG004260 ............... ............... ............... .............-- --------------- --------------- -K............. ............... ............... ............... 117
Psa_09bG004270 --------------- -------------.S .SMVPPLGA---.F. LSARLQPS.KVTI.L C.............. ............... ............... ............... ............... ............... 119
Psa_09bG004290 --------------- --------------- --------------- --------------- --------------- --------------- --------------- --------------- --------------- ---------------
Psa_09bG004310 --------------- --------------- --------------- --------------- -.............. ............... ............... ............... ............... ...............  89
Psa_09bG004320 ............... ............... ............... ............... ............... ............... ............... ............... ............... ............... 150
Psa_09cG004680 --------------- --------------- --------------- --------------- -----------MP.S SI.KTLERYQRCTFT PQTIHPNDHETLNWY KELSK.K.KYES.QR SQ-----------SI TKT--LERYQRCTFT  66
Psa_09cG004690 -----MD-G.WLGPS IASCCTIIFFPSSD. LLWW.TSVVR.ALPF PVGRRHPLRPLIIHV .L..--FHVKVYGAA LRRHFLVEC.--... ............... .............S. ............... ............... 140
Psa_09cG004710 ............... ............... ............... ............... ............... ............... ............... .............S. ............... ............... 150

Consensus     EL-KKKERQLGDINK QLKHKLGADGGSMRA LQGSWRPASGANIDT FRNHSSNMDTEPTLQ IGRYNQYVPSEATIP RNGGAGNTFMPGWGA VLV 242
Psa_09aG004230 ..-............ ............... ............... ............... ............... ............... .-- 101
Psa_09aG004260 ..-............ ............... ............... ............... ............... ............... .-- 207
Psa_09bG004270 ..-............ ............... ............... ............... ............... ............... .-- 209
Psa_09bG004290 --------------- ------------... ............... ............... ............... ............... I..  66
Psa_09bG004310 ..-............ ............... ............... ............... ............... ............... .-- 179
Psa_09bG004320 ..-............ ............... ............... ............... ............... ............... .-- 240
Psa_09cG004680 PQTIHPNDHETLNWY KELS..KEKY----- --------------- -------------ES LQ.SQRLTKF.G.ST TGKTT---------- --- 113
Psa_09cG004690 ..-............ ............... ............... ............... ............... ............... .-- 230
Psa_09cG004710 ..-............ ............... .....Q......... ............... ............... ............... .-- 240
```

**B**

| | $mS_{S7}$ | $mS_{S9}$ | $mS_{S11}$ | $lS_{S7}$ | $lS_{S9}$ | $lS_{S11}$ | $P_{S7}$ | $P_{S9}$ | $P_{S11}$ | $L_{S7}$ | $L_{S9}$ | $L_{S11}$ | $St_{S7}$ | $St_{S9}$ | $St_{S11}$ | $O_{S7}$ | $O_{S9}$ | $O_{S11}$ | $G_{S7}$ | $G_{S9}$ | $G_{S11}$ |
|---|---|---|---|---|---|---|---|---|---|---|---|---|---|---|---|---|---|---|---|---|---|
| Psa_09aG004230 | 0.00 | 0.00 | 0.00 | 11.58 | 19.01 | 10.91 | 0.00 | 0.00 | 0.00 | 106.33 | 116.50 | 78.48 | 0.00 | 0.47 | 1.29 | 8.13 | 5.45 | 5.08 | 5.09 | 4.81 | 6.44 |
| Psa_09aG004260 | 0.01 | 0.00 | 0.01 | 10.40 | 14.81 | 10.65 | 0.00 | 0.00 | 0.00 | 80.66 | 81.63 | 68.71 | 0.05 | 0.07 | 0.45 | 6.99 | 3.87 | 4.04 | 3.31 | 4.42 | 5.07 |
| Psa_09bG004270 | 0.01 | 0.00 | 0.00 | 14.96 | 22.07 | 14.48 | 0.00 | 0.02 | 0.02 | 129.30 | 136.50 | 115.60 | 0.00 | 0.27 | 0.55 | 10.71 | 5.82 | 5.99 | 5.21 | 6.90 | 9.06 |
| Psa_09bG004290 | 0.00 | 0.00 | 0.00 | 0.19 | 0.50 | 0.47 | 0.00 | 0.00 | 0.00 | 3.23 | 3.62 | 2.96 | 0.00 | 0.00 | 0.12 | 0.05 | 0.03 | 0.31 | 0.23 | 0.24 | 0.06 |
| Psa_09bG004310 | 0.00 | 0.00 | 0.00 | 5.65 | 7.03 | 5.70 | 0.00 | 0.02 | 0.00 | 44.56 | 50.19 | 45.05 | 0.00 | 0.02 | 0.02 | 4.18 | 1.82 | 1.84 | 1.73 | 2.46 | 3.16 |
| Psa_09bG004320 | 0.00 | 0.06 | 0.02 | 24.28 | 30.37 | 26.97 | 0.44 | 0.00 | 0.04 | 129.02 | 143.99 | 141.20 | 0.07 | 0.14 | 0.13 | 13.29 | 7.88 | 7.58 | 6.12 | 8.35 | 7.84 |
| Psa_09cG004680 | 0.00 | 0.00 | 0.00 | 0.00 | 0.00 | 0.00 | 0.00 | 0.00 | 0.00 | 0.00 | 0.00 | 0.00 | 0.00 | 0.00 | 0.00 | 0.00 | 0.00 | 0.00 | 0.00 | 0.00 | 0.00 |
| Psa_09cG004690 | 0.00 | 0.01 | 0.01 | 3.01 | 4.92 | 3.23 | 0.00 | 0.11 | 0.03 | 44.96 | 43.42 | 33.81 | 0.00 | 0.07 | 0.00 | 2.46 | 2.08 | 2.05 | 1.54 | 1.46 | 2.23 |
| Psa_09cG004710 | 0.03 | 0.00 | 0.00 | 18.85 | 27.03 | 18.41 | 0.00 | 0.27 | 0.13 | 143.62 | 169.31 | 175.69 | 0.00 | 0.07 | 0.00 | 14.67 | 11.70 | 13.51 | 8.93 | 9.98 | 9.31 |

**Fig. S6 Protein sequence alignment (A) and expression (B) of *PsAGL6-2* HGs.** HGs with truncated sequences were indicated with green shadow.

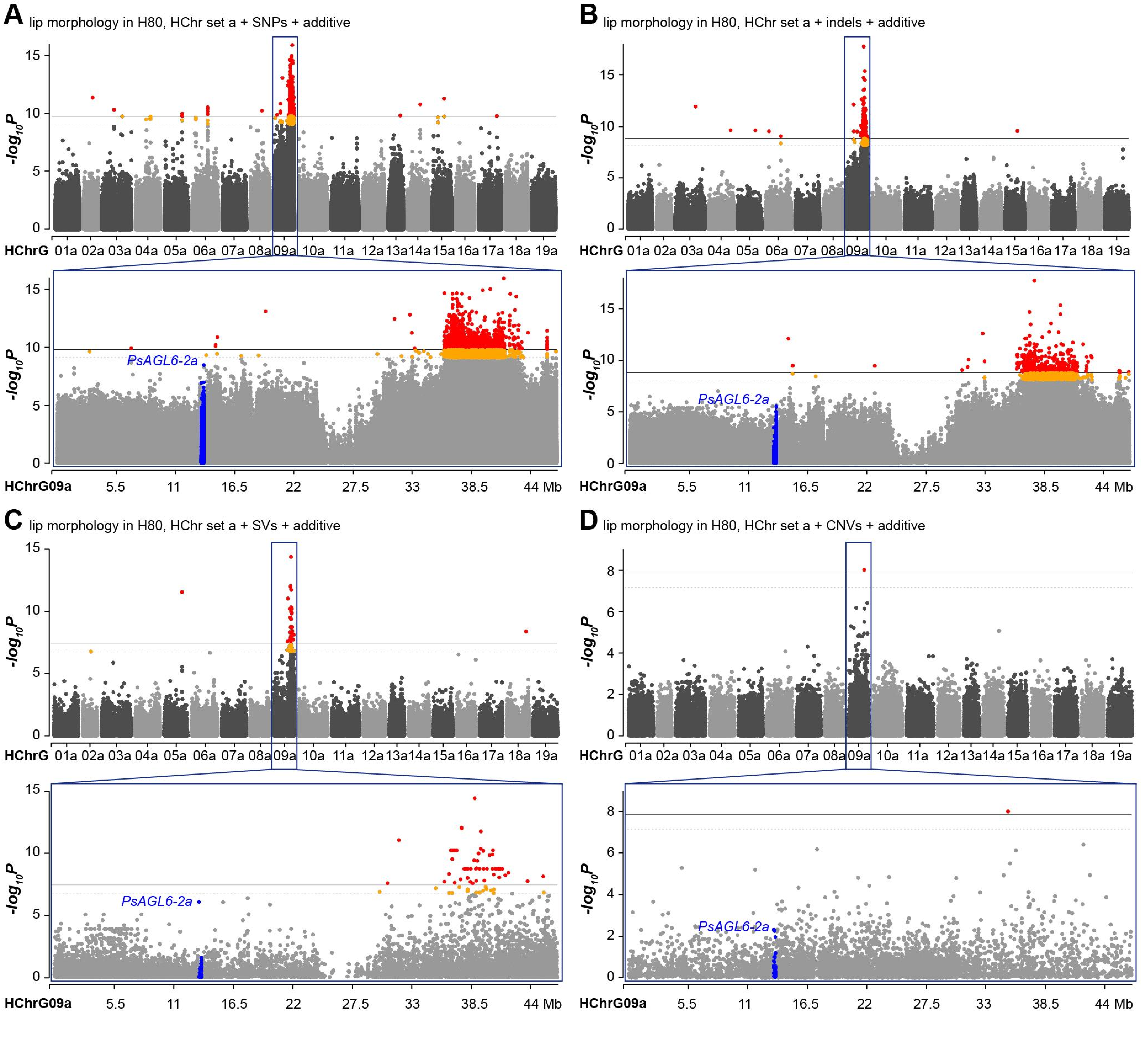


**Fig. S7 Manhattan plot for GWAS of lip morphology in the H80 population using HChr set a as the reference.** (**A-D**) GWAS results when SNPs, indels, SVs and CNVs were used, respectively, with the additive marker-effect model. Horizontal solid line and dashed line indicate $-\log_{10}$ (0.01/variant counts) and $-\log_{10}$ (0.05/variant counts) (**Methods**), respectively; red and orange dots denote variants significantly associated with the lip morphology variation; blue dots represent variants located within or adjacent (±2 kb) to the tandem duplicate cluster of *PsAGL6-2a* genes.

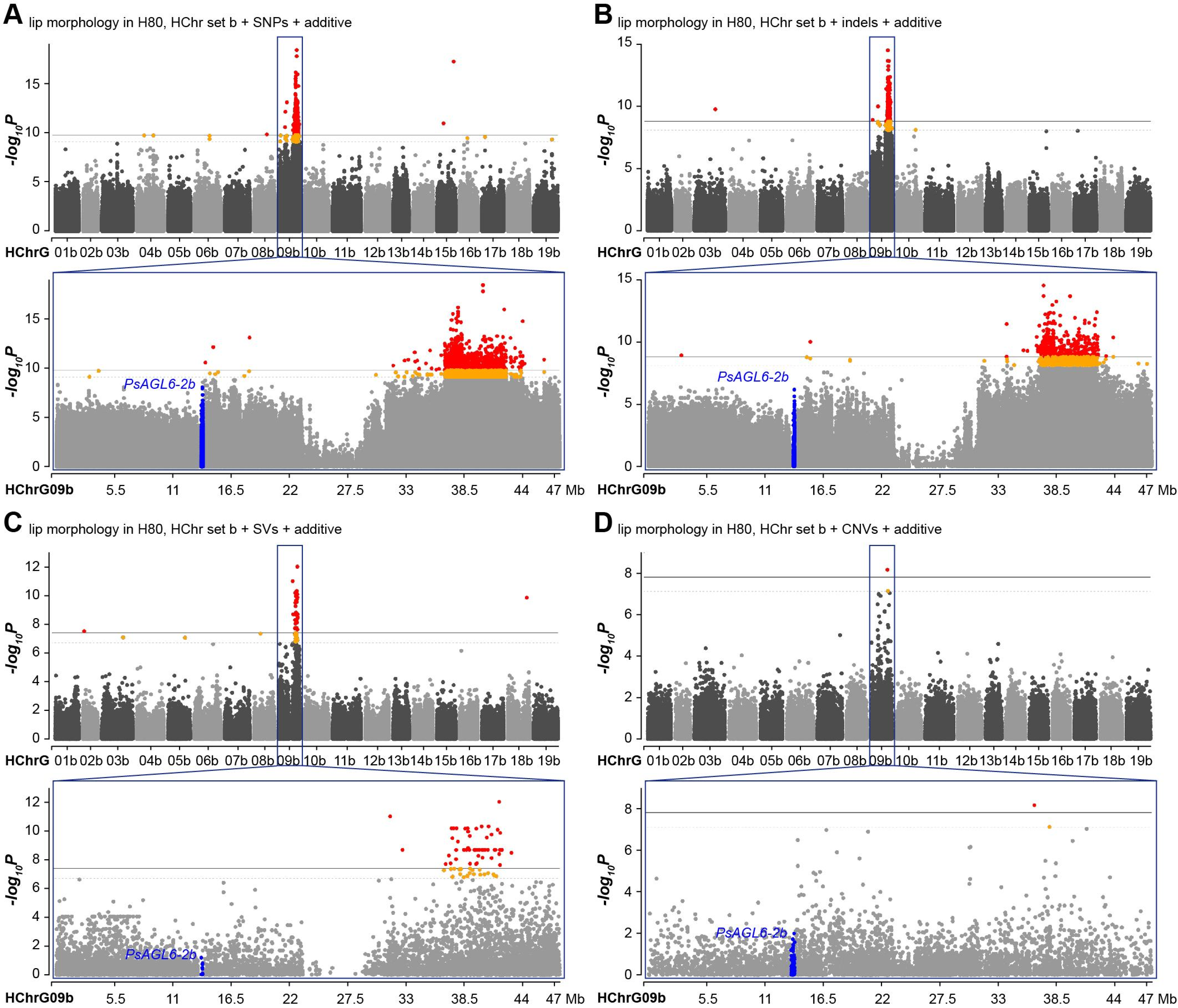


**Fig. S8 Manhattan plot for GWAS of lip morphology in H80 using HChr set b as the reference.** (**A-D**) GWAS results when SNPs, indels, SVs and CNVs were used, respectively, with the additive marker-effect model. Red and orange dots denote variants significantly associated with the lip morphology variation; blue dots represent variants located within or adjacent (±2 kb) to the tandem duplicate cluster of *PsAGL6-2b* genes.

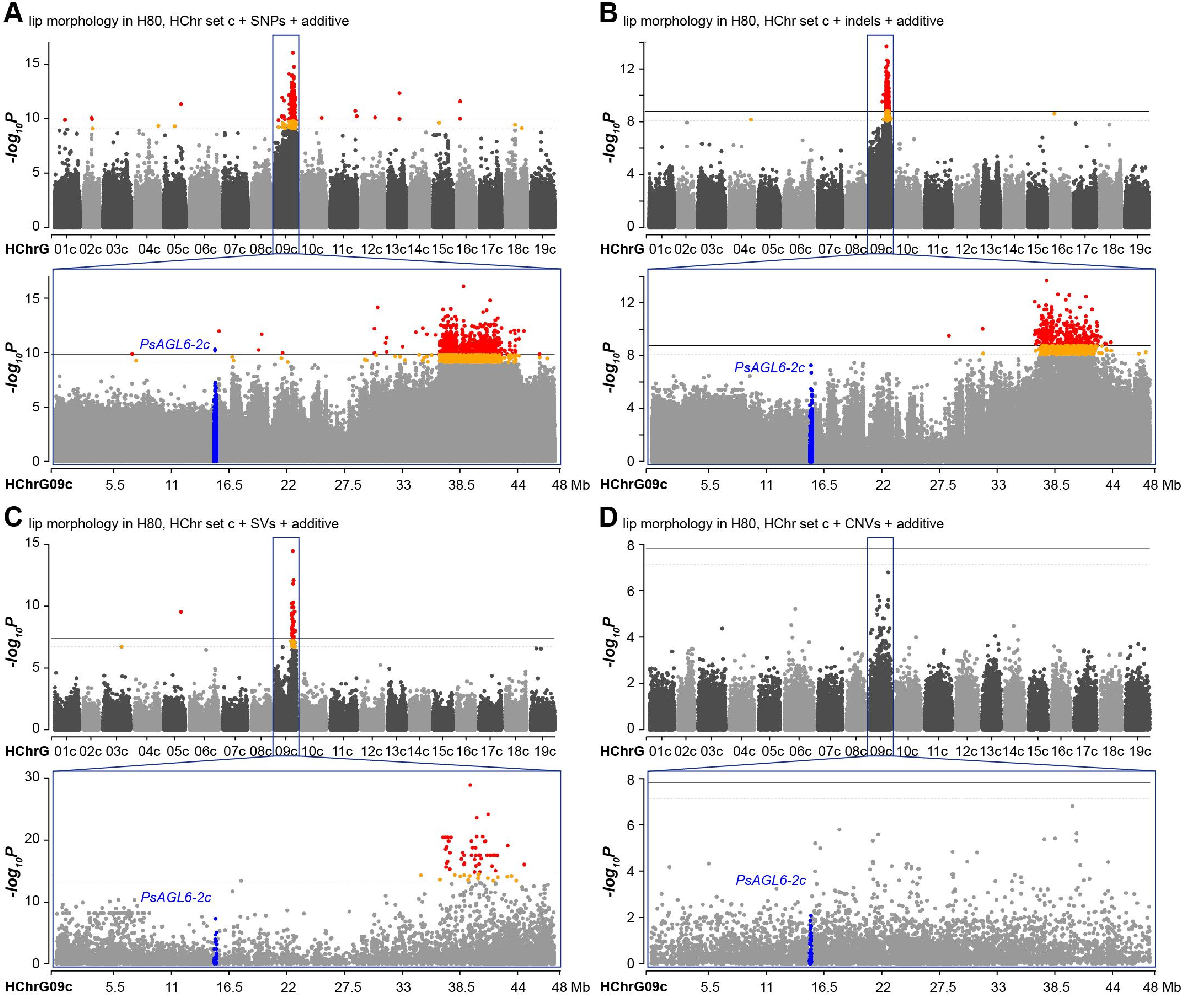


**Fig. S9 Manhattan plot for GWAS of lip morphology in H80 using HChr set c as the reference.** (**A-D**) GWAS results when SNPs, indels, SVs and CNVs were used, respectively, with the additive marker-effect model. Red and orange dots denote variants significantly associated with the lip morphology variation; blue dots represent variants located within or adjacent (±2 kb) to the tandem duplicate cluster of *PsAGL6-2c* genes.

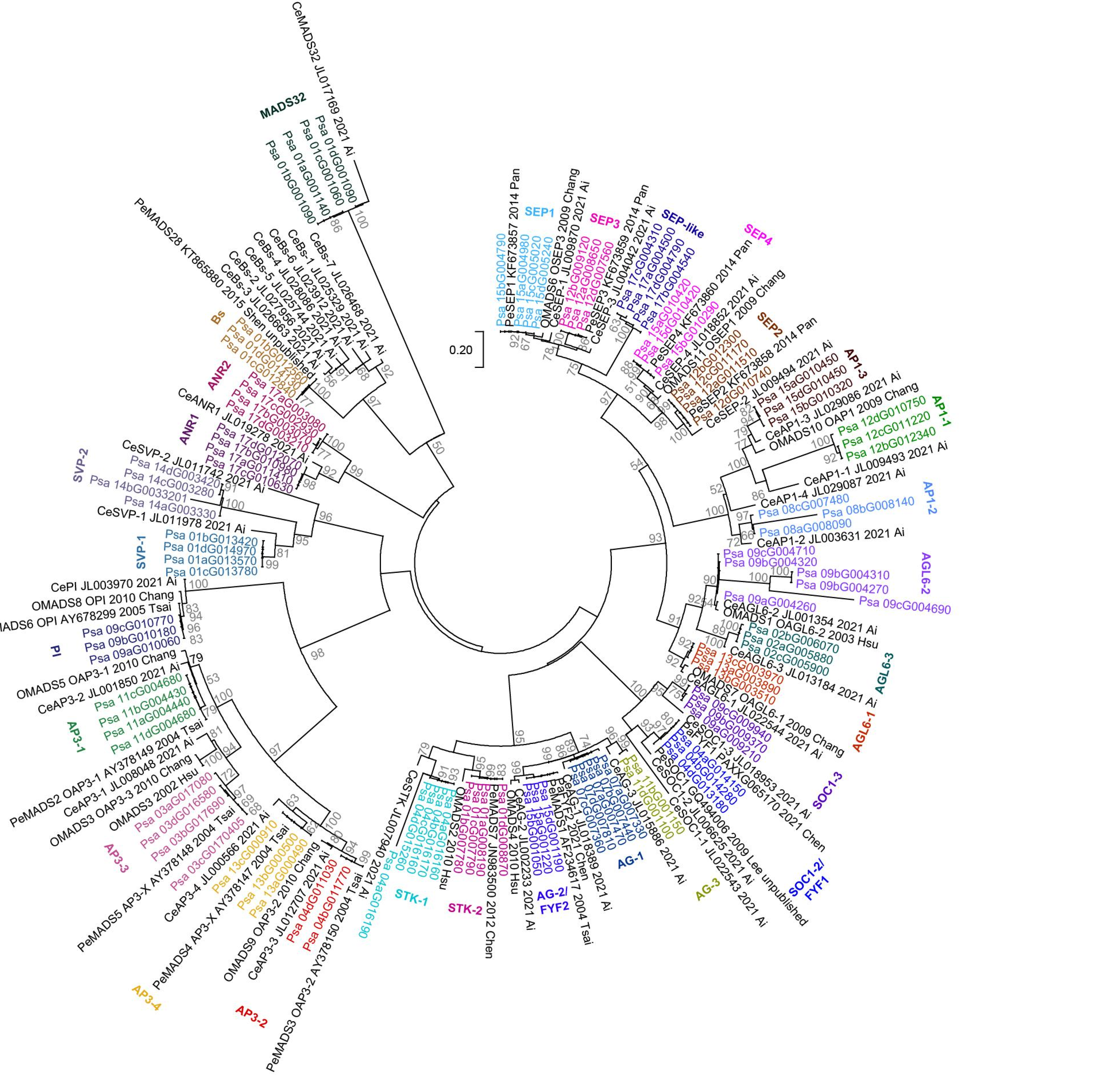


**Fig. S10 Gene tree for MIKC-type MADS-box genes in orchids.** NCBI accession ID (such as JL017169) was listed between gene name (such as *CeMADS32*) and the reference (such as 2021 Ai).

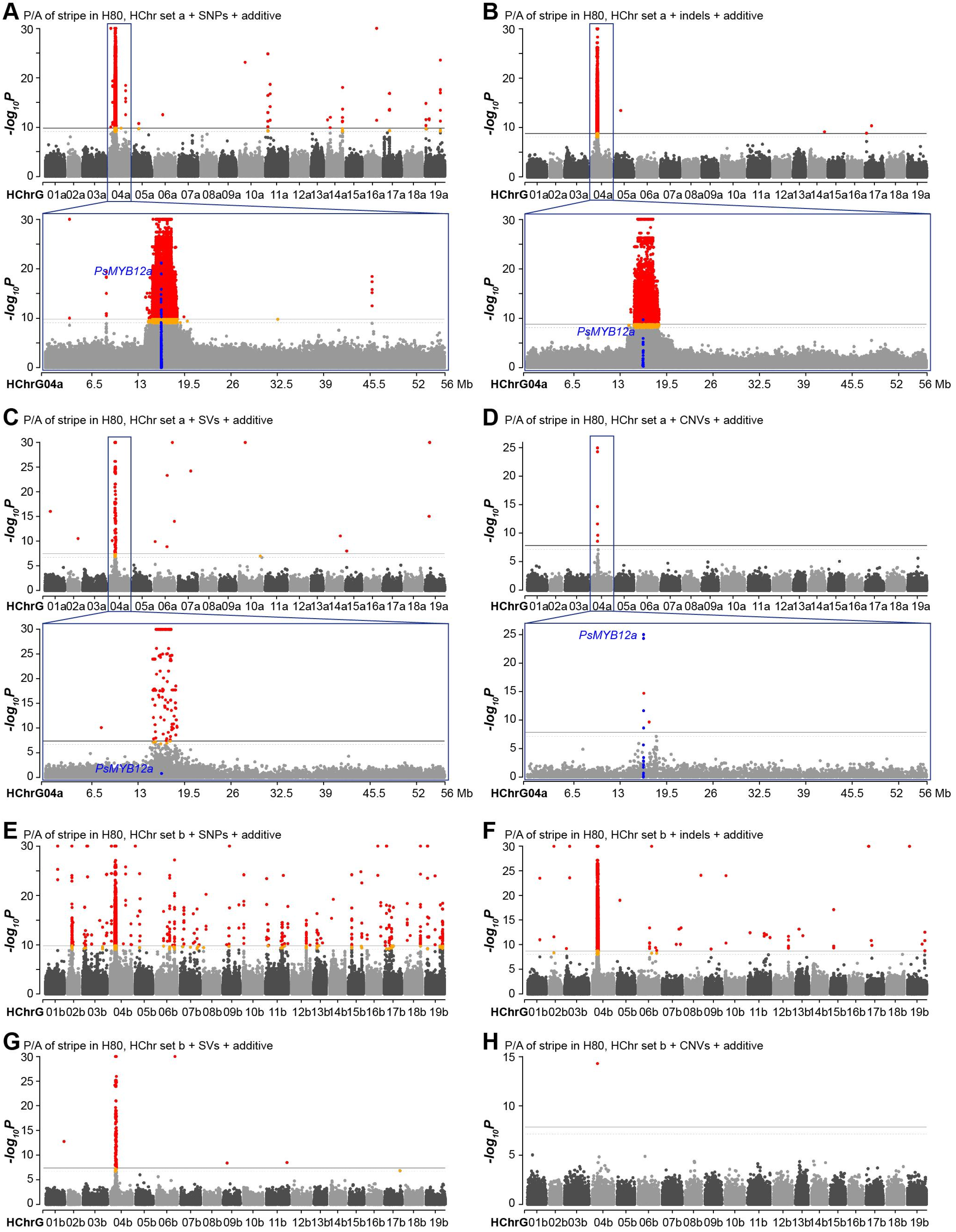


**Fig. S11 Manhattan plot for GWAS of stripe P/A in H80 using the additive marker-effect model.** (**A-D**) GWAS results when SNPs, indels, SVs and CNVs that were called by referring to HChr set a were used, respectively. (**E-H**) GWAS results when SNPs, indels, SVs and CNVs that were called by referring to HChr set b were used. Red and orange dots denote variants significantly associated with the P/A variation of stripe; blue dots represent variants located within or adjacent (±2 kb) to the tandem duplicate cluster of *PsMYB12a* genes.

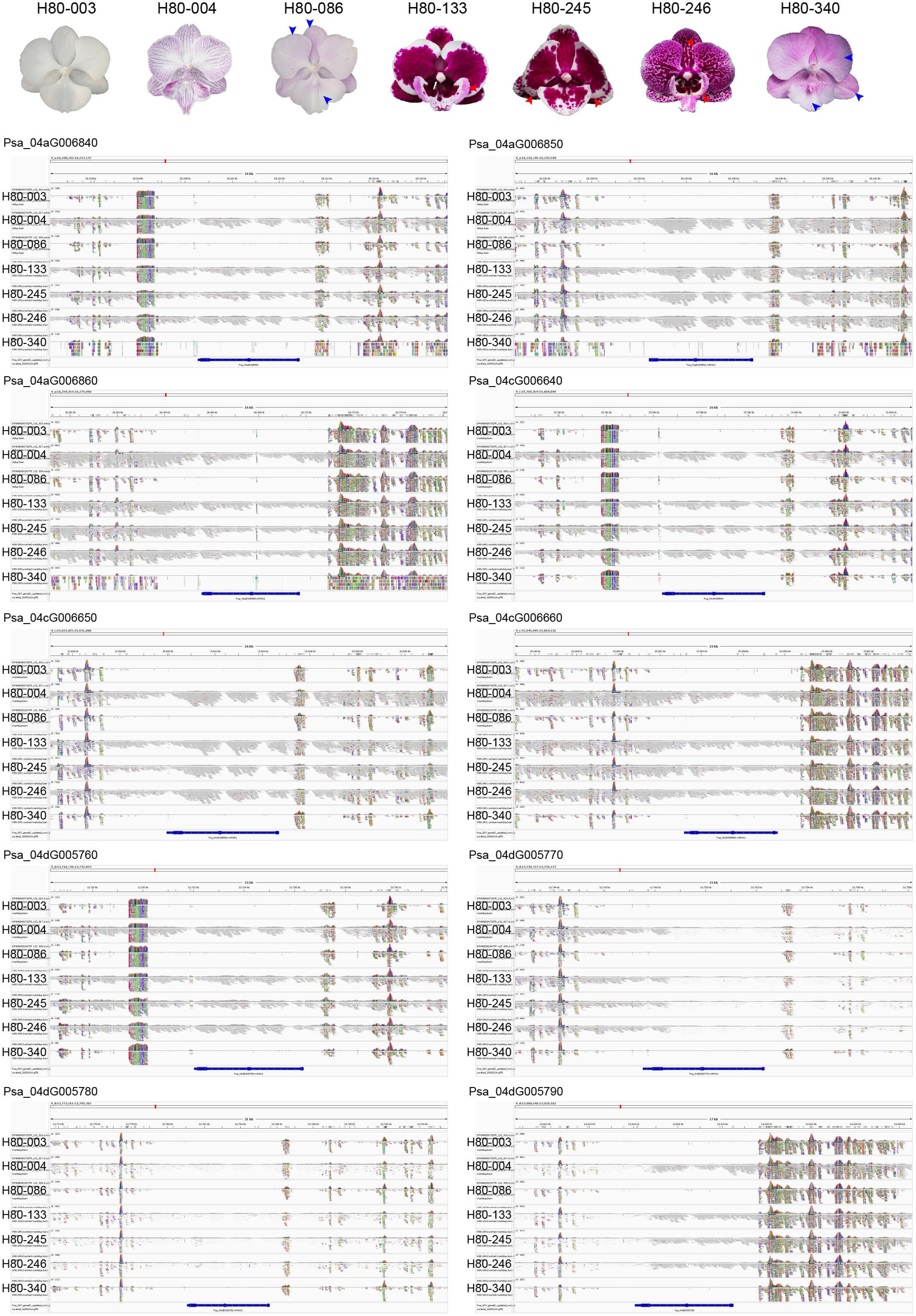


**Fig. S12 IGV images showing reads of seven H80 individuals with or without stripes that were mapped onto the *PsMYB12* gene-containing regions.** Red arrowheads indicate stripes, while blue ones indicate venation-associated shadow in individuals which had no intact reads mapped onto *PsMYB12* genes.

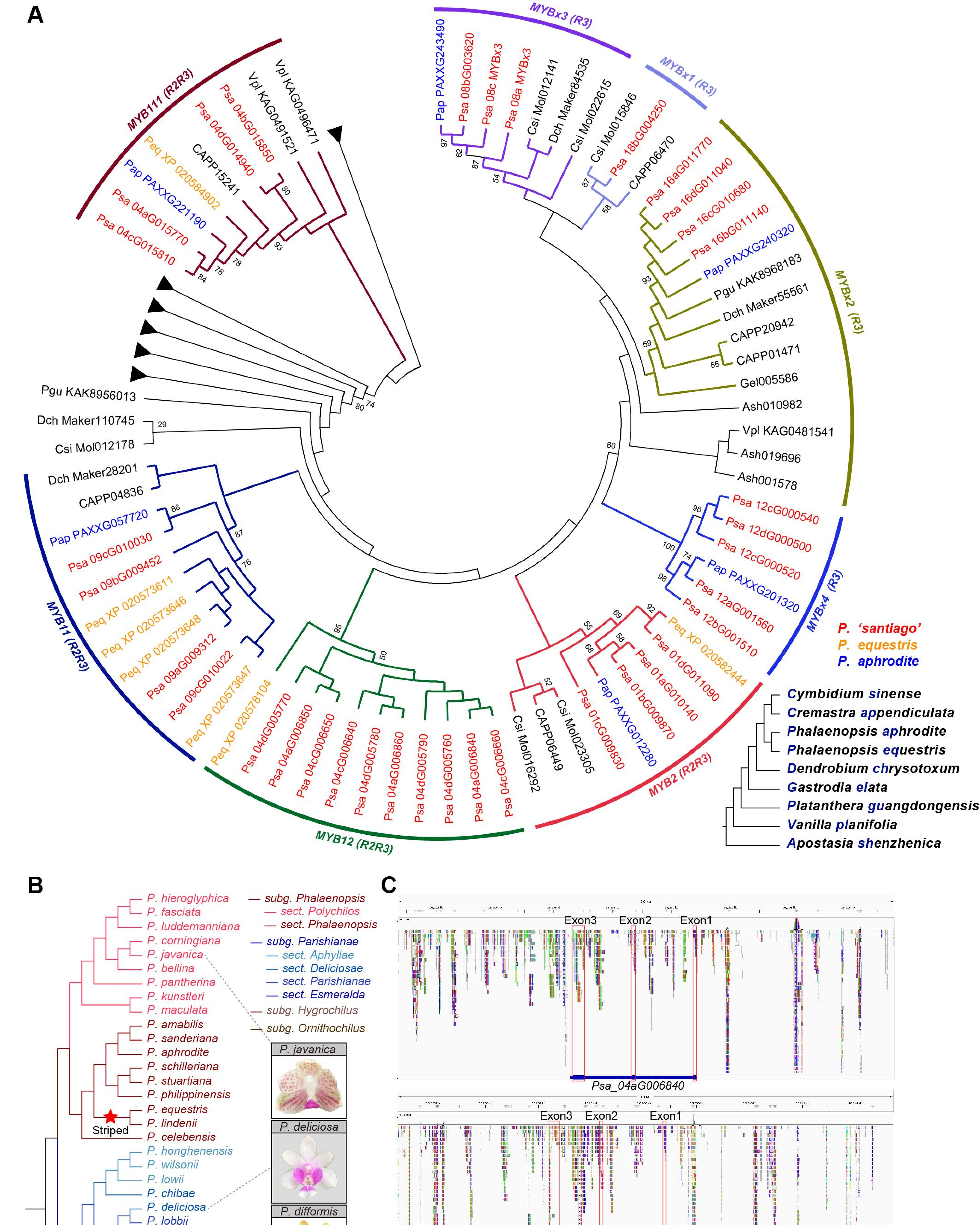


**Fig. S13 Evolution of color patterning-related *MYB* genes.** (**A**) Gene tree of *MYB* genes from orchids with genome assemblies. (**B**) Evolutionary history of stripes on perianths in the genus *Phalaneopsis*. The phylogeny of *Phalaenopsis* species were modified from our previous review paper (Wang *et al.*, 2025). The red star indicates the origination of stripes in the most recent common ancestor of *P. equestris* and *P. lindenii*. Three flower images show the venation-associated distribution of spots in other *Phalaenopsis* species. (**C**) Integrative Genomics Viewer images showing reads of *Vanda* 'Pakchong Blue' that were mapped onto genic regions of two *PsMYB12* genes as examples.

**A** ***PsMYB2* protein sequences**

```
                                  R2 -W-(X19)-W-(X19)-W-                                                  R3 -F/I-(X18)-W-(X18)-W-
Consensus      MGRNPSCSKEGLNKG AWTAVEDKLLIKYVN AHGEGKWTTVPHMAG LKRSGKSCRLRWLNY LRPNVKRGNFSEEEI DLIIRLHKLLGNRWS LIAGRIPGRTDNEIK NYWNTTLAKKTTSQL PSQKPPASNQALPSP SSSPSPQSTNNKTLN 150
Psa_01aG010140 ............... ............... ............... ............... ............... ............... ............... ............... ............... ............... 150
Psa_01bG009870 ............... ............... ............... ............... ............... ............... ............... ............... ............... ............... 150
Psa_01cG009830 --------------- --------------- --------------- --------------- -------MEIGNQRK FKYNQ.VSF.EI... ............... ............... ............... ............... 83
Psa_01dG011090 ............... ............... ............I.. ............... ............... ............... ............... ..............H ............... ...L........... 150

Consensus      RPKALRCSQLGFLVG EPNATVASGLPNQNT KASTIAEEELLQEEE ENVLWNCRFDDDNAV FADEDLMEFNGMQDF ESWMLNDDGCYLPDD ENINWLTSLFDMEGE LLKQF*                                           261
Psa_01aG010140 ............... ............... ............... ..............T. ............... ............... ............... ......                                           261
Psa_01bG009870 ............... ..I............ ............... ............... ............... ............... ............... ......                                           261
Psa_01cG009830 ............... ............... ............... ............... ............... ............... ............... ......                                           194
Psa_01dG011090 ............... ..T.A...V...... ............... ...V...S..EY... ............... ............... ............... ......                                           261
```

**B** ***PsMYB2* expression**

| | $mS_{S7}$ | $mS_{S9}$ | $mS_{S11}$ | $lS_{S7}$ | $lS_{S9}$ | $lS_{S11}$ | $P_{S7}$ | $P_{S9}$ | $P_{S11}$ | $L_{S7}$ | $L_{S9}$ | $L_{S11}$ | $St_{S7}$ | $St_{S9}$ | $St_{S11}$ | $O_{S7}$ | $O_{S9}$ | $O_{S11}$ | $G_{S7}$ | $G_{S9}$ | $G_{S11}$ |
|---|---|---|---|---|---|---|---|---|---|---|---|---|---|---|---|---|---|---|---|---|---|
| Psa_01aG010140 | 7.36 | 10.07 | 1.35 | 7.12 | 5.60 | 0.73 | 1.92 | 1.41 | 1.45 | 0.05 | 0.03 | 0.06 | 0.05 | 0.00 | 0.05 | 16.20 | 17.97 | 10.28 | 0.01 | 0.02 | 0.00 |
| Psa_01bG009870 | 32.00 | 37.80 | 2.58 | 24.82 | 22.93 | 1.38 | 8.00 | 9.14 | 5.94 | 0.43 | 0.32 | 0.50 | 0.02 | 0.00 | 0.00 | 32.38 | 42.76 | 26.95 | 0.08 | 0.06 | 0.00 |
| Psa_01cG009830 | 2.16 | 2.90 | 0.15 | 1.84 | 1.38 | 0.08 | 0.27 | 0.72 | 0.49 | 0.00 | 0.00 | 0.00 | 0.00 | 0.06 | 0.00 | 3.45 | 4.31 | 2.35 | 0.01 | 0.00 | 0.00 |
| Psa_01dG011090 | 9.98 | 7.71 | 0.02 | 7.52 | 0.75 | 0.02 | 2.34 | 0.54 | 0.00 | 0.18 | 0.01 | 0.17 | 0.00 | 0.00 | 0.00 | 9.11 | 8.94 | 3.14 | 0.19 | 0.24 | 0.05 |

| | $L_{003}$ | $L_{067}$ | $L_{064}$ | $L_{072}$ | $L_{014}$ | $L_{017}$ | $L_{077}$ | $L_{070}$ | $L_{128}$ | $L_{094}$ | $L_{324}$ |
|---|---|---|---|---|---|---|---|---|---|---|---|
| Psa_01aG010140 | 0 | 4.92 | 6.24 | 0.40 | 0.59 | 1.33 | 0.33 | 0 | 1.62 | 0 | 0.03 |
| Psa_01bG009870 | 0 | 9.73 | 16.93 | 0.33 | 0.83 | 2.01 | 0.33 | 0.05 | 2.67 | 0.05 | 0 |
| Psa_01cG009830 | 0 | 5.95 | 9.17 | 0.59 | 0.89 | 1.18 | 0.27 | 0 | 1.00 | 0.06 | 0 |
| Psa_01dG011090 | 0 | 0.04 | 0.04 | 0 | 0 | 0 | 0 | 0 | 0 | 0 | 0 |

**C** ***PsMYB11* protein sequences**

```
Consensus      MKQKAKYDEDGVNKG AWTAEEDELLASYVN AVGEGHWTSVPKKAG LNRRGKSCRLRWLNY LRPNIKRGNISVEEE ELIIRLHKLLGNRWS LIAGRLPGRTDNEIK NYWNTTLSKKIQTKK FTINMPNIKAWKPKS NPLETKITSSISYSI 150
Psa_09aG009312 ............... ............... ............... ............... ............... ............... ............... ............... ............... ............... 150
Psa_09bG009452 ............... ............... ............... ............... ............... ............... ............... ............... ............... ............... 150
Psa_09cG010022 ............... ............... ............... ............... ............... ............... ............... ............... ............... ......T........ 150
Psa_09cG010030 ...R..N.....R.. ............C.. T....N..L.M.... ...G........... ............... ............... ............... ..............N ............... ............... 150

Consensus      QTTELKCTKASYPLH XLPSTSAVIQXQQIS QQNFQEELMEEKKDK VLDWDSFSFDDEMWI AGLEFGGEINVVDDY DNDLFCSLFDQNL*                                                                  239
Psa_09aG009312 ............... A.........S.... ............... ............... ............... ..............                                                                  239
Psa_09bG009452 ............... A.........S.... ............... ............... ............... ..............                                                                  239
Psa_09cG010022 ............... V.........T.... ............... ............... ............... ..........E...                                                                  239
Psa_09cG010030 ............... V.....G...T.... ............... ............... ..........A.... ....V.....A...                                                                  239
```

**D** ***PsMYB11* upstream 1 Kb sequences**

```
Consensus      CTATGATAATGAAGT AAACATAGCATTTGA AAAAA-GATACACCG AGGTAAGGTYTTAAT GATATTGTAAGTAGG GTCGTAAASGGACGA GCTCAGGCTTGGCTT GTGTACGGC---TTG TTTGAAATTAATTAT GTTCGAAATTGGCTT 146
Psa_09aG009312 ............... ............... .....-......... ............... ............... ............... ............... .........---... ............... ............... 146
Psa_09bG009452 ............... ............... .....-......... ............... ............... ............... ............... .........---... ............... ............... 146
Psa_09cG010022 ............... ............... .....A....A..T. ............... ............... ...........G... ............... .........---... ............... ............... 147
Psa_09cG010030 ............... ........TG...C. .....-..C..G.-A ............... ...........C... .C.......A..A.. ...........AT.. ....G....GGC... ............... ...T.........C 148

Consensus      TTGTTTGAAATTATT TGTGATCAATGTTTT CGTGTTTGATTTGAT TTATATATTCTTGTT CGTGTTTGGCTTATG TACAACTTGAAATTA GCTTGTCTAAATAAA TTAAAATAATAGATG TGGGGGAAAAAAATA GAACCATATGTATCG 296
Psa_09aG009312 ............... ............... ............... ............... ............... ............... ............... ............... ............... ............... 296
Psa_09bG009452 ............... ............... ............... ............... ............... ............... ............... ............... ............... ............... 296
Psa_09cG010022 ............... ............... ............... ............... ............... ............... ............... ............... ..........-.... ............... 296
Psa_09cG010030 A.............. .....A..G.....C .CCA.....C..... ...C..G..T..... ...........CGC. .T.T........... ......A...T.... ...G..A...TT..T ....-----.TGT.G ..TT..A..C.T.T. 293

Consensus      CATGAGTTGTGATGA TTTAAACTTATATAA ATATAACATGATCGA ATAAAAAAATGGTTTG TGTCAAGCTTGTTTA TAATTTGAATTTAAA AAATTTATTTGTG-- TTTGACTCGTTTATT TAAYGAATAATTCAA ACAATGAATCTAACA 444
Psa_09aG009312 ............... ............... ............... ............... ............... ............... .............-- ............... ............... ............... 444
Psa_09bG009452 ............... ............... ............... ............... ............... ............... .............-- ............... ............... ............... 444
Psa_09cG010022 ..C........C... ............... .............A. ......T........ .T..C.......... ............... .........C...-- ............... ............... ............... 444
Psa_09cG010030 ..---A..A.A..A. .---..TAA...G.. TC....G..C.CTCG .GTT.TG...A...A AAATT.-TA.AAAC. ...CAC..TCCA.C. ...GG.GACAAA.AA .C.A..AT.AGGTG. .C.....A...GACT TTTT..G.C---.GC 433

Consensus      CGAGCTATTCATGAA AAATGAGATTTTTTT TTTTTTTNNNTTGAC AGCCATGCTACTAAC AATAATAATATATTT GTATACAAACCACCT AACATTTTTGTATTT ACGTCTCAAACATTT TTTATCTATTTATTT TTTTTNATATTATTT 594
Psa_09aG009312 ............... ............... ............... ............... ............... ............... ............... ............... ............... ............... 594
Psa_09bG009452 ............... ............... ............... ............... ............... ............... ............... ............... ............... ............... 594
Psa_09cG010022 ............... ............... .....A.---..... ............... ............... ............... ............... ............... ............... .....-......... 590
Psa_09cG010030 ..TA...C.A..-.. T..CA.T..AA.A.A ...---------..G .TAA.CTT.GA..T. --...AGGACC...C .C...T...T..... ..TT........... ....A..G.....C. .....T..C...... ....----...TA.A 567

Consensus      TATATTACACAAAAA TTGTAATTATGTTAA AATTTTGGAAAGTGA AATGCCATCACGTTG ACGGGAACATGGCTA TATGTTAAGTTTTAA GTTTTGGTGTCAGGT ATGTWGGTACGTAGG GGGGCASCGGCCATG AGCTTTGTAGTATCT 744
Psa_09aG009312 ............... ............... ............... ............... ............... ............... ............... ............... ............... ............... 744
Psa_09bG009452 ............... ............... ............... ............... ............... ............... ............... ............... ............... ............... 744
Psa_09cG010022 ............... ............... ............... ............... ............... ............... ............... ............... ............... ........G...... 740
Psa_09cG010030 ...GCCT.T...TT. ..----....A...C .--.AGAA....... G...TAGA.G.ACGC .T..AT.ATGA.G.. .GC..--------.. .............AC ..TG.A........T ..A.--....G.... .....GACT..G.T. 701

Consensus      GTTTAATTAGCCGCA CATGCCTCATCCGAC GGTCAAGAAGAACCG AAACGG----CAGCT T-ATAAACCTACTAG TTAGGAGCAGAAAGG TCGTTCTGCAGTCTC TAAACTTAGTACTCT TAAATAACTCCCTTA TATTCTCCCACTTAT 889
Psa_09aG009312 ............... ............... ............... ......----..... .-............. ............... ............... ............... ............... ............... 889
Psa_09bG009452 ............... ............... ............... ......----..... .-............. ............... ............... ............... ............... ............... 889
Psa_09cG010022 ............... .....G......... ............... ......----..... .-............. ............... ............... ............... ............... ............... 885
Psa_09cG010030 .CAC.G.A...T... ...........T... ..........T..T. ...A.AGGGC...T. CG..........G.. C..........C.A C.T.....TC.GT.. ....T.......G.A ............A.C ..AC..T.T....G. 851

Consensus      TTTCATATA--AAAA TTCAGGAGACATATC TATCTAAGTTTATTT ATTTAAGCGGCCTGA GGTGAGAAGMAGCAG CAGCAGCAGCAACAG TAGCAGCGGCACAAG AAGCAGAG                                        1000
Psa_09aG009312 .........--.... ............... ............... ............... ............... ............... ............... ........                                        1000
Psa_09bG009452 .........--.... ............... ............... ............... ............... ............... ............... ........                                        1000
Psa_09cG010022 .........--.... ............... ............... ............... ............... ............... ............... ........                                        996
Psa_09cG010030 ..AT.....GA.... A..T..G.....G.. .............C. ........A....A. .T............. ........A..G..A C.A....A.T.G... ....G...                                        964
```

**E** ***PsMYB11* expression**

| | $mS_{S7}$ | $mS_{S9}$ | $mS_{S11}$ | $lS_{S7}$ | $lS_{S9}$ | $lS_{S11}$ | $P_{S7}$ | $P_{S9}$ | $P_{S11}$ | $L_{S7}$ | $L_{S9}$ | $L_{S11}$ | $St_{S7}$ | $St_{S9}$ | $St_{S11}$ | $O_{S7}$ | $O_{S9}$ | $O_{S11}$ | $G_{S7}$ | $G_{S9}$ | $G_{S11}$ |
|---|---|---|---|---|---|---|---|---|---|---|---|---|---|---|---|---|---|---|---|---|---|
| Psa_09aG009312 | 0.53 | 0.55 | 1.92 | 0.69 | 2.66 | 3.59 | 0.44 | 0.66 | 1.22 | 0.15 | 0.94 | 5.58 | 1.08 | 1.34 | 1.00 | 0.47 | 2.00 | 1.01 | 0.78 | 0.68 | 1.37 |
| Psa_09bG009452 | 0.31 | 0.57 | 2.19 | 0.79 | 3.32 | 3.43 | 0.38 | 0.38 | 1.31 | 0.17 | 0.89 | 5.31 | 1.16 | 1.46 | 1.40 | 0.47 | 1.60 | 1.14 | 0.97 | 0.70 | 1.72 |
| Psa_09cG010022 | 0.88 | 1.14 | 2.31 | 2.01 | 2.99 | 3.28 | 1.69 | 0.29 | 2.95 | 1.12 | 0.98 | 6.93 | 0.96 | 2.82 | 1.94 | 0.54 | 1.84 | 1.04 | 1.53 | 0.93 | 2.37 |
| Psa_09cG010030 | 0.01 | 0.03 | 0.14 | 0.05 | 1.33 | 3.69 | 0.00 | 0.04 | 0.10 | 0.19 | 0.96 | 41.42 | 0.00 | 0.00 | 0.00 | 0.00 | 0.17 | 0.35 | 0.00 | 0.00 | 5.57 |

| | $L_{003}$ | $L_{067}$ | $L_{064}$ | $L_{072}$ | $L_{014}$ | $L_{017}$ | $L_{077}$ | $L_{070}$ | $L_{128}$ | $L_{094}$ | $L_{324}$ |
|---|---|---|---|---|---|---|---|---|---|---|---|
| Psa_09aG009312 | 0.06 | 3.57 | 0.11 | 6.07 | 7.04 | 6.03 | 19.02 | 34.62 | 0.63 | 1.13 | 0.23 |
| Psa_09bG009452 | 0.06 | 3.24 | 0.17 | 5.49 | 7.52 | 6.08 | 19.49 | 33.58 | 0.27 | 0.82 | 0.05 |
| Psa_09cG010022 | 0.06 | 1.15 | 0.28 | 2.89 | 3.19 | 1.68 | 10.85 | 15.53 | 0.54 | 0.21 | 0.00 |
| Psa_09cG010030 | 1.60 | 2.19 | 1.34 | 10.79 | 10.58 | 5.79 | 34.41 | 98.91 | 0.27 | 1.75 | 0.00 |

**Fig. S14 Homoeologous gene variation in sequence and expression of *PsMYB2* and *PsMYB11*.** (**A**) Protein sequence alignment of *PsMYB2* HGs. (**B**) HG-specific expression of *PsMYB2* in *P*. sa floral organs (upper) and the lips of selected H80 individuals (lower), with values in the table indicating TPM values. (**C,D**) Protein sequence and promoter DNA sequence (1 Kb upstream) alignments of *PsMYB11* HGs, respectively. (**E**) HG-specific expression of *PsMYB11* in *P*. sa floral organs (upper) and the lips of selected H80 individuals (lower). The HG showing high sequence divergence with others was indicated using green shadow.

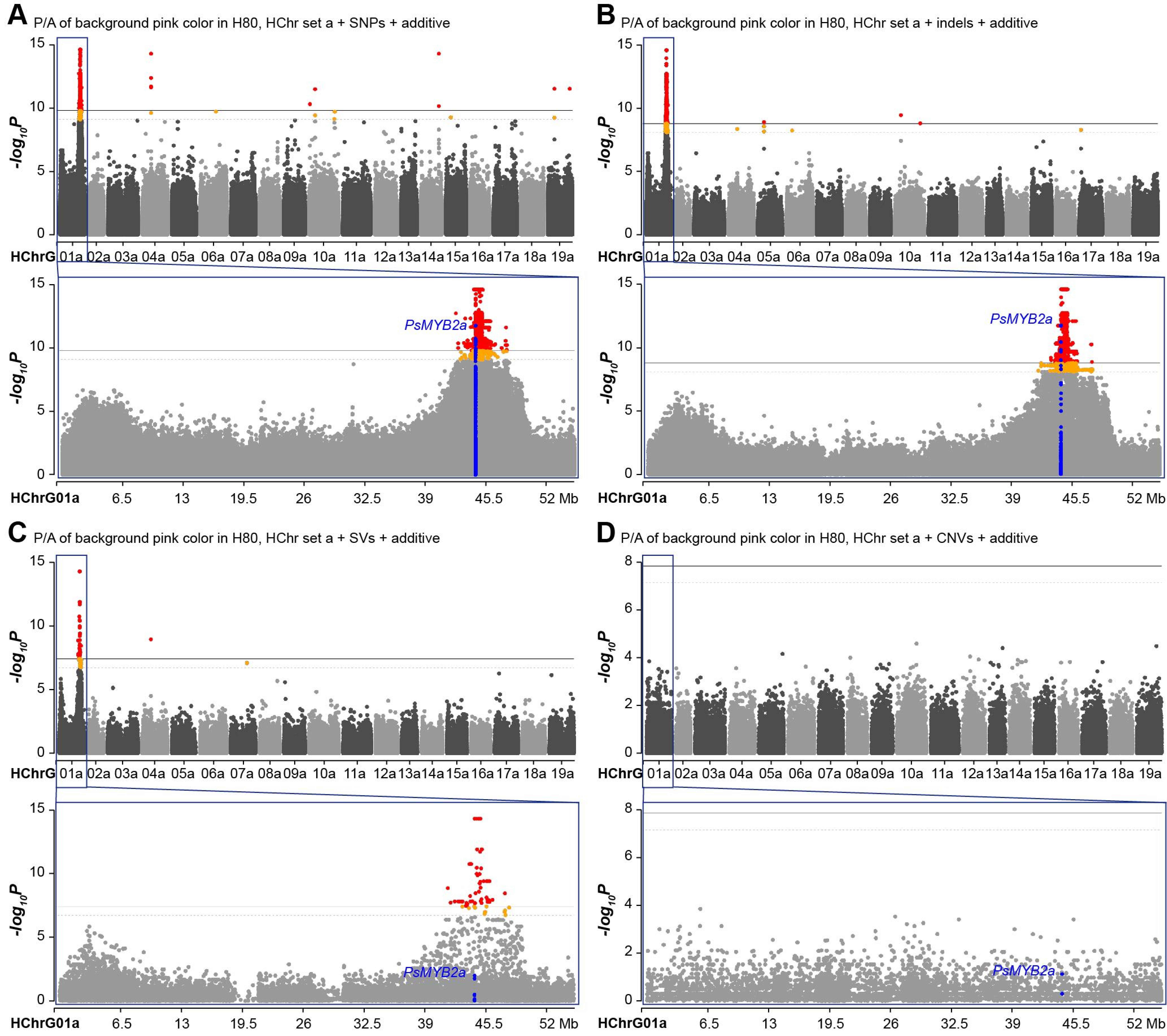


**Fig. S15 Manhattan plot for GWAS of background color in H80 using the additive marker-effect model.** (**A-D**) GWAS results when SNPs, indels, SVs and CNVs that were called by referring to HChr set a were used, respectively. Red and orange dots denote variants significantly associated with the P/A variation of pink color; blue dots represent variants located within or adjacent (±2 kb) to *PsMYB2a*.

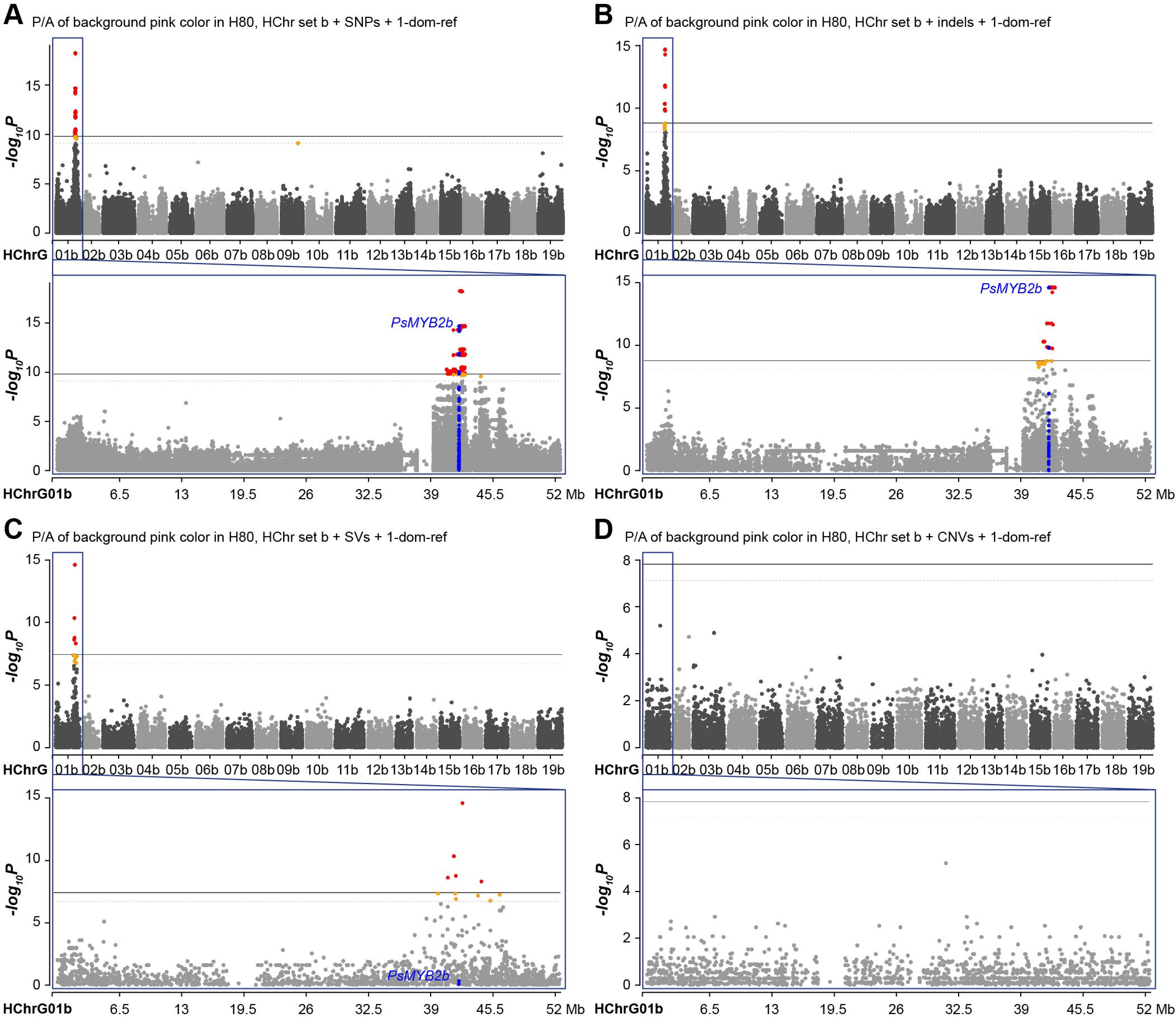


**Fig. S16 Manhattan plot for GWAS of background color in H80 using the 1-dom-ref marker-effect model.** (**A-D**) GWAS results when SNPs, indels, SVs and CNVs that were called by referring to HChr set b were used, respectively. Red dots denote variants significantly associated with the P/A variation of pink color; blue dots represent variants located within or adjacent (±2 kb) to *PsMYB2b*.

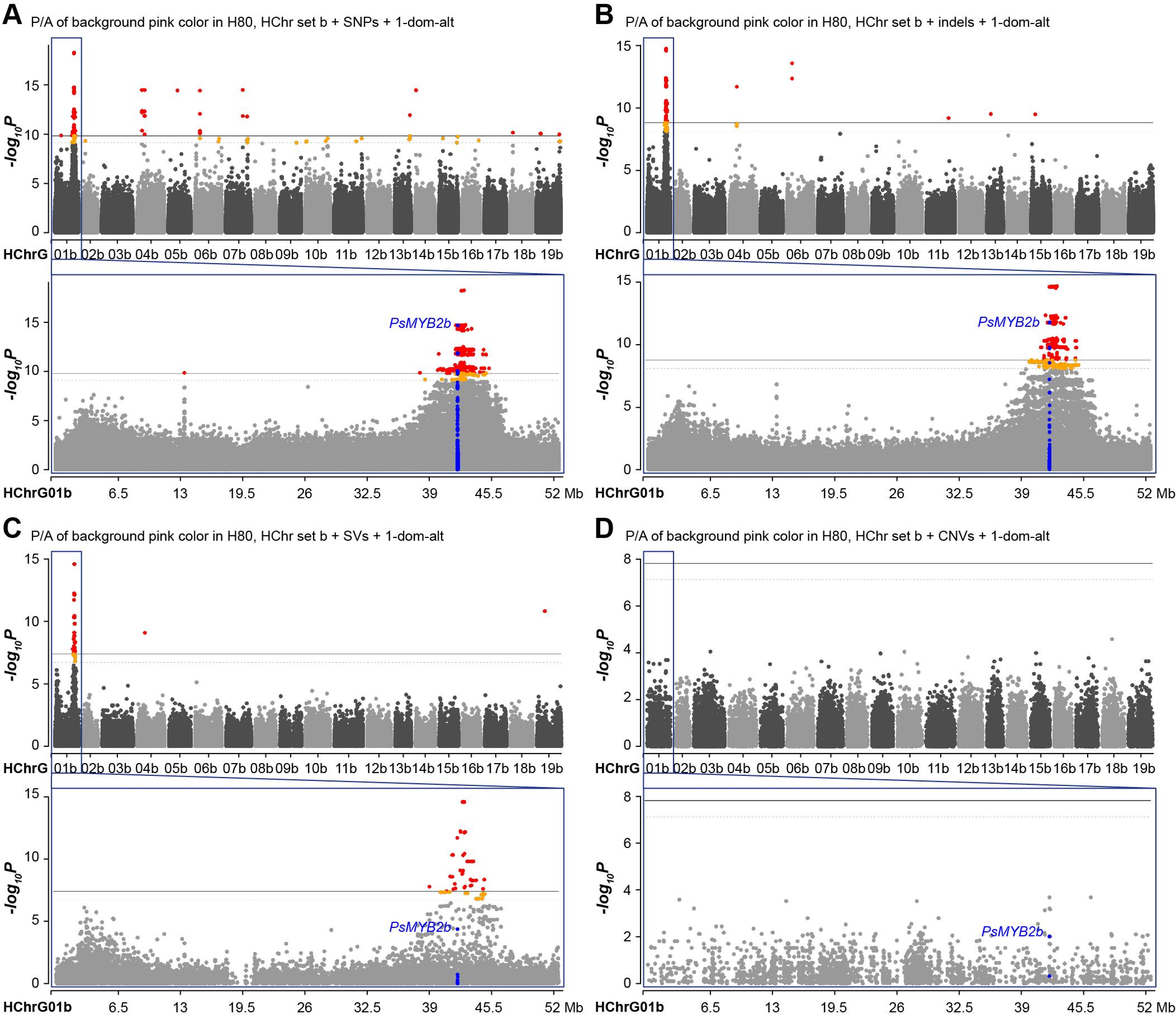


**Fig. S17 Manhattan plot for GWAS of background color in H80 using the 1-dom-alt marker-effect model.** (**A-D**) GWAS results when SNPs, indels, SVs and CNVs that were called by referring to HChr set b were used, respectively. Red and orange dots denote variants significantly associated with the P/A variation of pink color; blue dots represent variants located within or adjacent (±2 kb) to *PsMYB2b*.

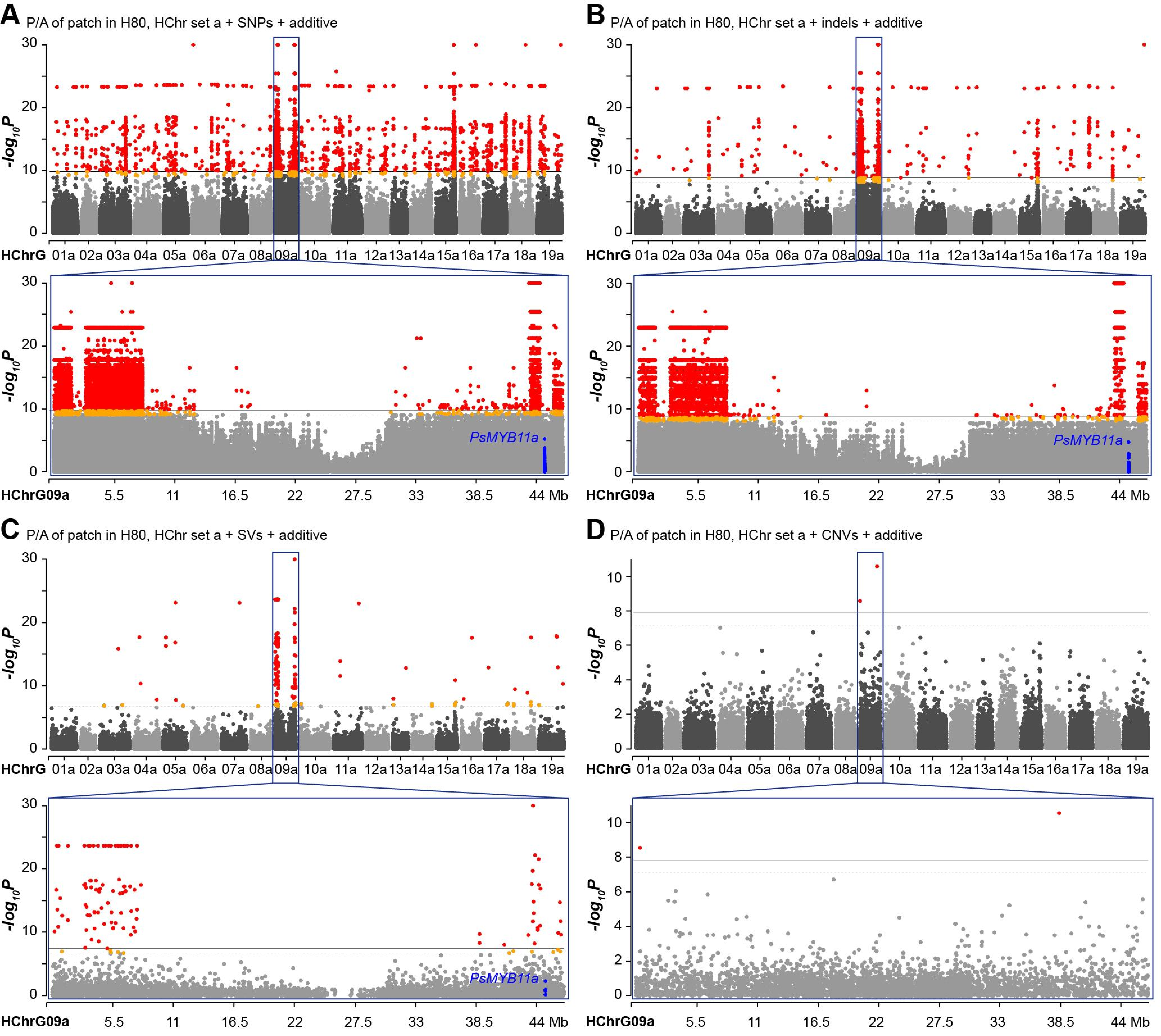


**Fig. S18 Manhattan plot for GWAS of patch P/A in H80 using HChr set a as the reference.** (**A-D**) GWAS results when SNPs, indels, SVs and CNVs were used, respectively, with the additive marker-effect model. Red and orange dots denote variants significantly associated with the P/A variation of patch; blue dots represent variants located within or adjacent (±2 kb) to *PsMYB11a*.

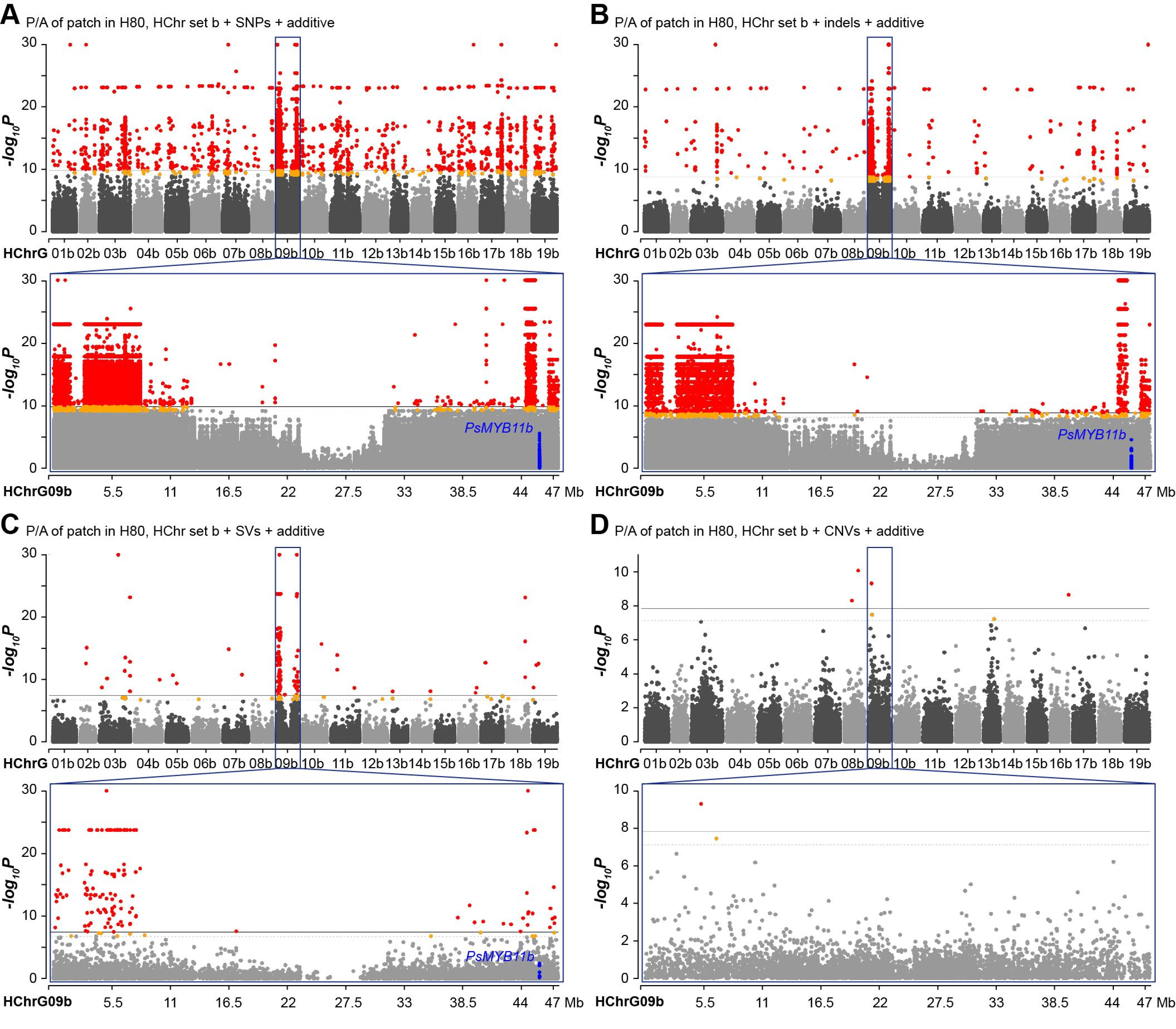


**Fig. S19 Manhattan plot for GWAS of patch P/A in H80 using HChr set b as the reference.** (**A-D**) GWAS results when SNPs, indels, SVs and CNVs were used, respectively, with the additive marker-effect model. Red and orange dots denote variants significantly associated with the P/A variation of patch; blue dots represent variants located within or adjacent (±2 kb) to *PsMYB11b*.

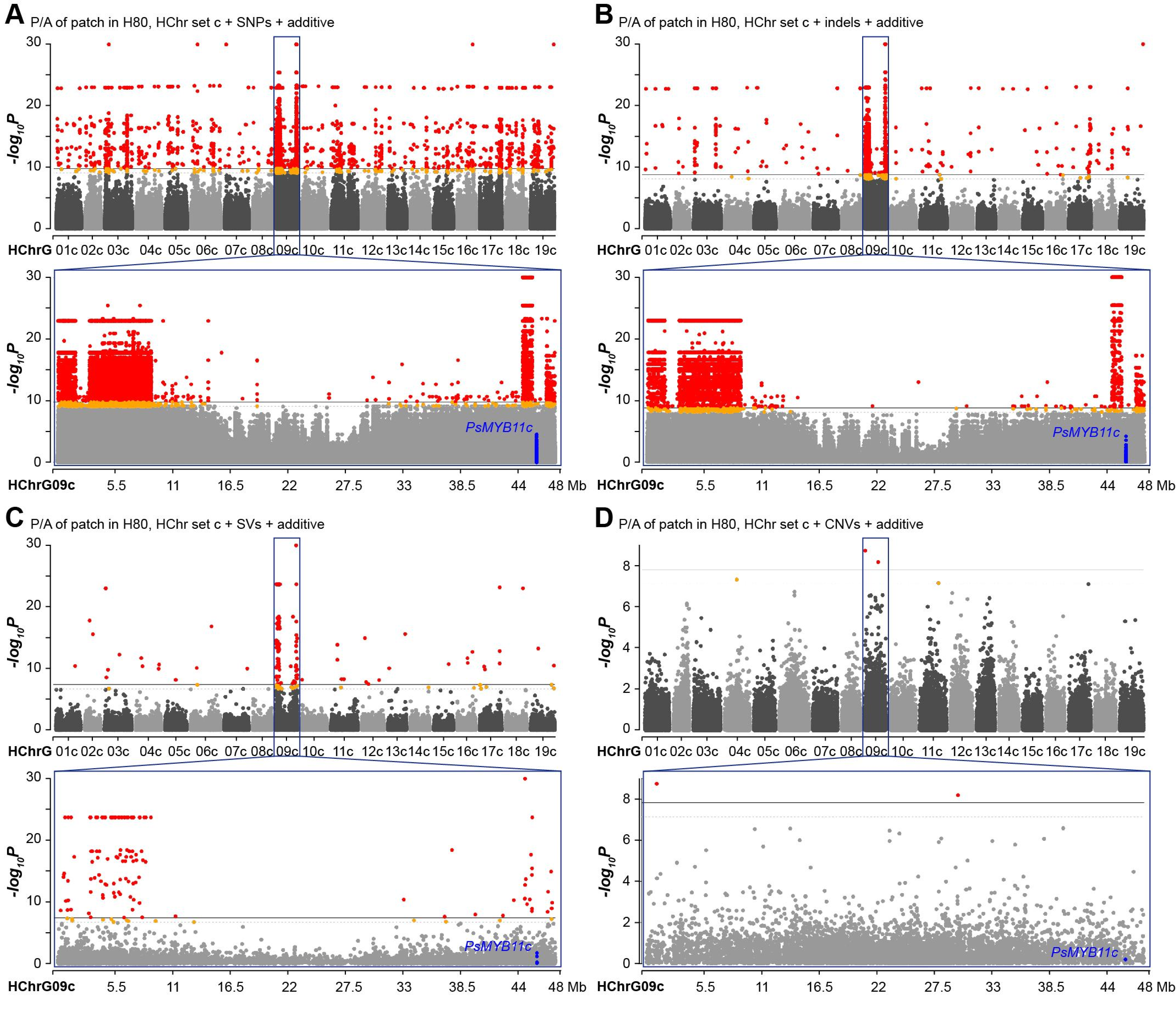


**Fig. S20 Manhattan plot for GWAS of patch P/A in H80 using HChr set c as the reference.** (**A-D**) GWAS results when SNPs, indels, SVs and CNVs were used, respectively, with the additive marker-effect model. Red and orange dots denote variants significantly associated with the P/A variation of patch; blue dots represent variants located within or adjacent (±2 kb) to the tandem duplicate cluster of *PsMYB11c* genes.

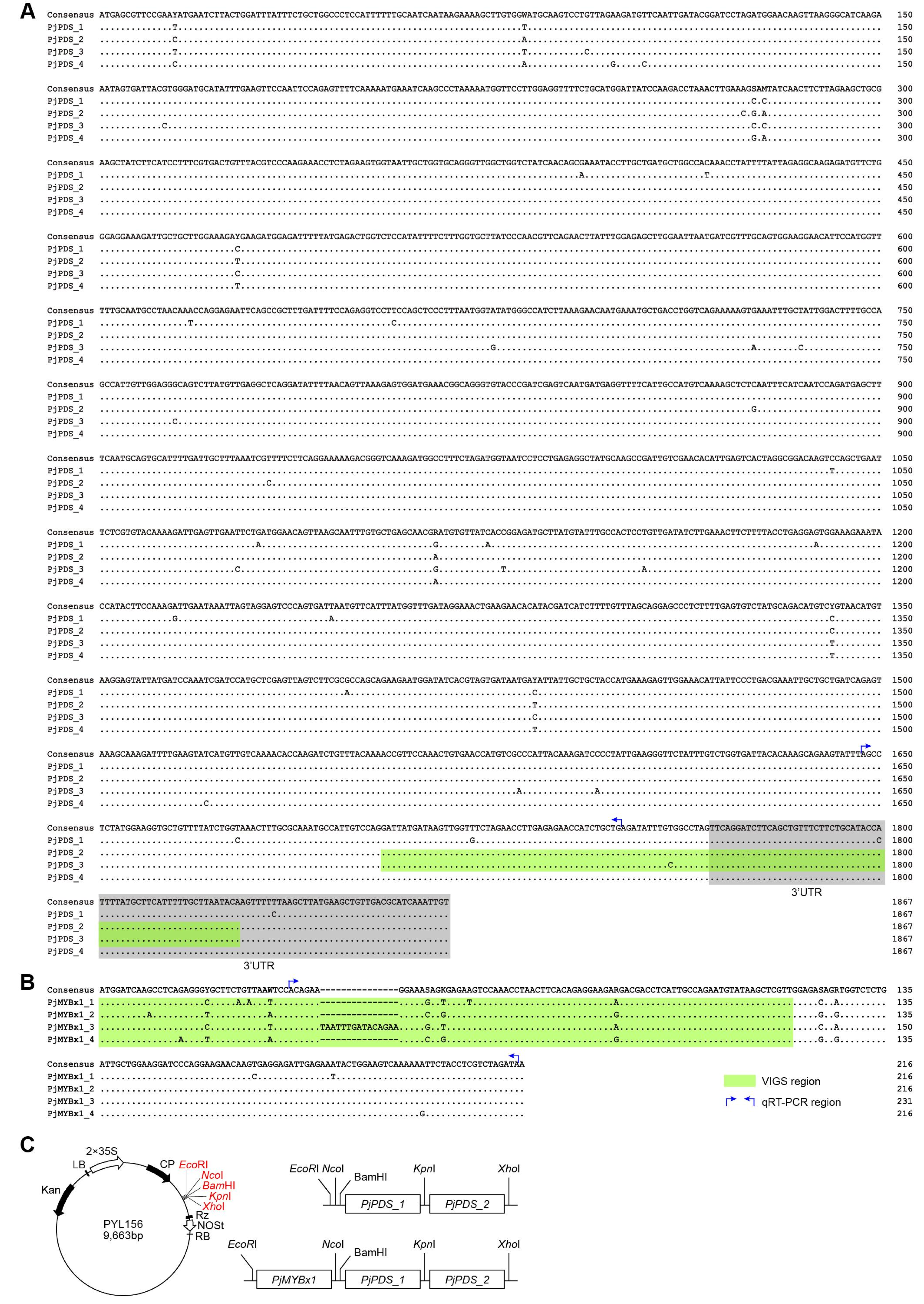


**Fig. S21 *PjPDS* and *PjMYBx1* sequences and the VIGS constructions.** (**A**) CDS and 3'UTR sequence alignment for four *PjPDS* HGs. (B) CDS sequence alignment for four *PjMYBx1* HGs. (C) Diagrams showing the structure of TRV2 plasmid *PYL156*, and the insertion locations of *PjPDS* and *PjMYBx1* sequences. *PjPDS_1* and *PjPDS_2* are sequences from two *PjPDS* HGs marked with green shadow.
